# Risk reshapes amygdala representation of choice

**DOI:** 10.1101/2025.10.06.680813

**Authors:** Patrick T. Piantadosi, Kendall M. Coden, Hyesun Choi, Sarah J. Perry, Marcelle Halfeld, Julia A. Schaffer, Jeff P. Goff, Nathen A. Spitz, Nicole R. Schwab, Rodrigo Sandon, Shanzeh Sadiq, Maddison M. Devine, Vincent D. Costa, Daniel da Silva, Andrew Holmes

## Abstract

Modifying behavior in response to changing environmental conditions is a crucial adaptive function. This capacity is exemplified when animals curtail pursuit of a valued outcome that risks being punished by aversive consequences, but the mediating brain mechanisms remain poorly understood. Here, using *in vivo* cellular-resolution calcium (Ca^2+^) imaging, optogenetics and chemogenetics, we show that risk of punishment dramatically alters animals’ choice between a large/risky and small- /safe reward and produces novel, causally necessary, patterns of activity in basolateral amygdala (BLA) neurons. We find that experience of a punished outcome generates a BLA representation that is selectively replayed when animals subsequently abort choice of the large/risky reward option. Additionally, we show that risk leads to the incorporation of newly encoding BLA neurons into the pre-choice representation, which predicts shifting away from the large/risky option. These findings reveal how dynamic reshaping of BLA representations underpins behavioral flexibility in the face of risk.

## 1 INTRODUCTION

Pursuing a goal often involves the risk of punishment, such as potentially encountering a predator while foraging, and learning to adjust behavior in response to such risks is highly adaptive (Jean-Richard-Dit-Bressel et al. 2018, Piantadosi et al. 2021). In humans, tolerance for risk varies across individuals and is aberrant in neuropsychiatric conditions. Alcohol and substance use disorders are defined in part by insensitivity to punishment risk (Ariesen et al. 2023, Brevers et al. 2014, Fishbein et al. 2005, Grant et al. 2000) (with rodent models of addiction characterized by similar alterations, see Marchant et al. 2013, Vanderschuren et al. 2017) whereas excessive risk aversion typifies mood and anxiety disorders (Giorgetta et al. 2012, Hockey et al. 2000, Mitte 2007, Stöber 1997, Yuen and Lee 2003). In rodents, punishment risk induces a state characterized by indecision and the exploration and exploitation of alternative sources of reward (if available) (Halladay et al. 2019, Hunt and Brady 1955, Simon et al. 2009).

Previous studies investigating the neural loci underlying behavioral adaptation under risk strongly implicate the amygdala. Humans with amygdala lesions perform poorly on decision-making tasks wherein gains are associated with risk of loss (Bechara et al. 1999, Brand et al. 2007, Weller et al. 2007). Likewise, rodents with lesion or inactivation of the basolateral amygdala (BLA) maintain reward-seeking despite punishment (Choi and Kim 2010, Ishikawa et al. 2020, Jean-Richard-Dit-Bressel and McNally 2015, Marchant et al. 2019, Orsini et al. 2015, Pelloux et al. 2013, Piantadosi et al. 2017, Verharen et al. 2019). Moreover, BLA neurons flexibly track changes in action value and encode positive and negative valence via subpopulations separable based on a multitude of factors, including activity profile, genetic identity, anatomical localization, and projection-target (Belova et al. 2007, Beyeler et al. 2016 2018, Burgos-Robles et al. 2017, Courtin et al. 2022, Kim et al. 2016, Kyriazi et al. 2018, Lim et al. 2024, Morse et al. 2020, Namburi et al. 2015, Paton et al. 2006, Piantadosi et al. 2024b, Shen et al. 2019, Tye and Janak 2007, Zhang et al. 2013 2021, Zhang and Li 2018). These findings illustrate that neurons in BLA are positioned to encode risk-related variables, in line with the BLA’s broader role in regulating behavioral responses to stimuli based on expected outcomes (Schoenbaum et al. 1998 1999, Saez et al. 2017, Bermudez and Schultz 2010, Balleine et al. 2003). Yet, key questions remain regarding how activity within BLA neuronal populations represent and regulate behavior when a rewarded action is subsequently punished.

Here, we address these questions by utilizing cellular-resolution Ca^2+^ imaging and complementary causal manipulations to examine how BLA neuronal activity encodes choice-related information as mice pursue a valued outcome that becomes associated with varying probabilities of punishment. We find that BLA representations of choice adapt to reflect risk and enable behavioral adaptations that are in part read-out through a projection to medial nucleus accumbens shell (NAcSh).

## 2 RESULTS

### Punishment risk shifts choice in a BLA-dependent manner

We trained food-restricted mice to initiate trials and choose between two spatially-separated square stimuli on a touchscreen: when touched, one stimulus produced a large volume ( 20 *µ*L, Large^Rew^) flavored milk reward and the possibility of a minor footshock punishment (0.1 mA), whereas the other stimulus produced a smaller volume ( 5 *µ*L, Small^Rew^) but was never punished (**Fig. 1a**). Animals were first given ∼13 reward-magnitude (RM) discrimination training sessions during which there was zero probability of punishment in order to engender reliable choice (>90%) of the Large^Rew^ option by the final (Late) RM session (**Extended Data Fig. 1a, Supplementary Video 1**). On the following session, mice underwent a risky-decision test (RDT, modified from Glover et al. 2020, Simon et al. 2009) comprising one block of safe trials followed by two blocks of ‘risky’ trials where the percentage of punished Large^Rew^ trials ascended in succession, first to 50% and then to 75% (Fig. 1b).

**FIGURE 1.**
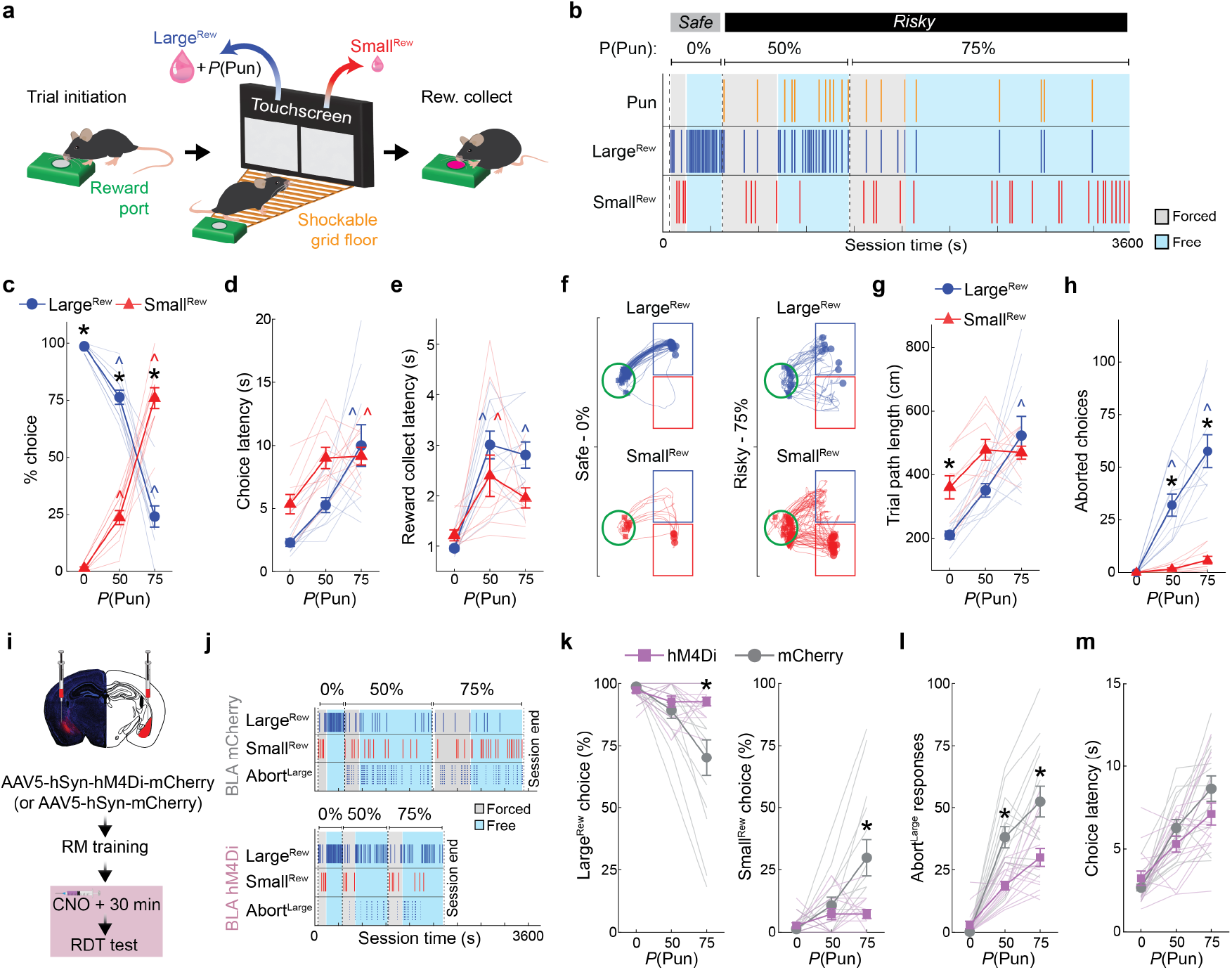
Choice behavior reflects the balance of reward value and punishment risk. (**a**) Cartoon depicting RDT task-structure sequence, left to right: head-entry in the reward port initiates a trial and displays a square visual stimulus within the touchscreen response window(s), touching a stimulus produces a Large^Rew^ ( 20 μL) and potential footshock punishment (P(Pun)) or a Small^Rew^ ( 5 μL), delivered in the reward port. (**b**) Raster plot from a representative RDT session showing footshocks received (Pun) and Large^Rew^ and Small^Rew^ responses across safe (0%) and risky (50 & 75%) trial blocks. Forced (stimulus on one screen only) and free (stimulus on each screen) trials shaded in grey and blue, respectively. (**c**) Mice (n=10, undergoing pan-BLA microendoscopic imaging) made fewer Large^Rew^ and more Small^Rew^ choices as punishment risk increased across blocks (Large^Rew^ vs Small^Rew^: p<0.001 for all blocks; within Trial Block effect for Large^Rew^ and Small^Rew^: all p<0.001; ANOVA interaction, F(2,18)=180.69, p<0.001). (**d**) Longer choice latencies on the riskiest versus the safe block for both choice-types (within Trial Block effect for Large^Rew^ and Small^Rew^: both p<0.001 for 75% risky block versus safe; ANOVA interaction, F(2,18)=4.37, p<0.05). (**e**) Longer reward collection latencies on risky versus safe blocks for both choice-types (within Trial Block effect for Large^Rew^: p<0.001 for both risky blocks versus safe block; within Trial Block effect for Small^Rew^: p<0.001 for the 50% risky block vs safe block; ANOVA interaction, F(2,18)=5.39, p<0.05). (**f**) Representative trial-wise locomotor paths on safe (left) and 75% risky (right) block in zones near the reward port zone (green circle) and the Large^Rew^ (blue square) and Small^Rew^ (red square) response-windows. (**g**) Longer locomotor paths for Small^Rew^ versus Large^Rew^ on the safe block (Large^Rew^ vs Small^Rew^: p<0.05 for safe block), and for Large^Rew^ on the riskiest block versus the safe block (within Trial Block effect for Large^Rew^: p<0.001 for 75% risky block versus safe; ANOVA interaction, F(2,18)=6.29, p<0.009). (**h**) More aborted choices directed at the Large^Rew^ versus Small^Rew^ response-window during risky trial-blocks (Large^Rew^ vs Small^Rew^: p<0.001 for risky blocks) and more Large^Rew^ aborts as compared to the safe block (within Trial Block effect for Large^Rew^: p<0.001 for the 50% and 75% risky blocks versus safe; ANOVA interaction, F(2,18)=24.75, p<0.001). (**i**) Representative hM4Di-mCherry expression in BLA (top); experimental timeline for chemogenetic manipulation (bottom). (**j**) Rasters as in **b**, depicting behavior from representative mCherry control (top) and hM4Di mice (bottom). (**k**) More Large^Rew^ (left) and fewer Small^Rew^ (right) choices on the riskiest trial-block in hM4Di mice (n=15) versus mCherry controls (n=14) (hM4Di vs mCherry: p<0.001 on the 75% risky block for Large^Rew^ and Small^Rew^; ANOVA interactions, F(2,54)=9.05, p<0.005). (**l**) Fewer Large^Rew^ aborts in hM4Di versus controls on both risky blocks (hM4Di vs mCherry: p<0.001 for risky blocks; ANOVA interaction, F(2,54)=8.41, p<0.005). (**m**) No group difference in choice latency. All multiple comparisons (Tukey’s post-hoc tests) were conducted following significant Trial Block (0, 50, 75%) x Choice Type (Large^Rew &^ Small^Rew^) or Trial Block x Treatment (hM4Di & mCherry) ANOVA interactions. For full reporting of statistics, see **Supplementary Table 1**). *p<0.05 versus other choice type (**c,g,h**) or mCherry control (**k,l**). ^p<0.05 versus safe block, color corresponding to choice type (**c-e, g,h**). Data mean ± SEM. Mouse (**a**, doi.org/10.5281/zenodo.3925913) reward water drop (**a**, doi.org/10.5281/zenodo.3925935), Hamilton Syringe (**i**, doi.org/10.5281/zenodo.7679042), and Syringe (**i**, doi.org/10.5281/zenodo.4152947) graphics adapted from scidraw.io.

The introduction of punishment during the RDT profoundly affected reward-seeking actions. We found that mice made fewer Large^Rew^ and more Small^Rew^ choices (**Fig. 1b,c**) and spent less time near the Large^Rew^ option on the risky, relative to safe, RDT blocks (**Extended Data Fig. 1b,c**). Behavior during risky blocks also slowed, as reflected in longer choice and reward-collection latencies and lengthier trial initiation-to-choice locomotor paths (**Fig. 1d-g**). Furthermore, punishment risk increased the propensity for choices to be aborted, i.e., mice approached the response-window and then retracted without executing a choice (**Fig. 1h, Supplementary Video 2**). Aborts occurred almost exclusively on approaches to the Large^Rew^, but not the Small^Rew^, response-window on risky blocks, suggesting that this behavior reflected the approach-avoid conflict associated with higher value, but now risky, option. Neither mouse weight nor footshock sensitivity were strongly correlated with Large^Rew^ preference on the RDT (**Extended Data Fig. 1d-f**), whereas motivation (assessed on a progressive ratio task) tended to be positively correlated with this RDT measure (**Extended Data Fig. 1g-i**).

On establishing the behavioral profile of mice in the RDT task, we used chemogenetic inhibition to assess the causal contribution of BLA to performance. To do so, we virally expressed the inhibitory designer receptor exclusively activated by designer drug (DREADD), hM4Di (or an mCherry control), in BLA neurons and systemically administered the DREADD ligand, clozapine N-oxide (CNO; 3 mg/kg), prior to RDT (**Fig. 1i,j, Extended Data Fig. 2a,b**). We found that hM4Di-expressing mice maintained greater Large^Rew^ preference during the riskiest trial-block than mCherry controls (**Fig. 1j,k**). Risk-related increases in aborted responses directed at the Large^Rew^ response-window (Abort^Large^) were also comparatively less numerous in the hM4Di group than controls, whereas choice latencies did not differ (**Fig. 1l,m**). By contrast, both groups exhibited comparable RDT performance when given vehicle instead of CNO (**Extended Data. Fig. 2c-e**). CNO-treated hM4Di-expressing mice reacted to the footshock-punisher less strongly than did controls during the RDT (Corder et al. 2019), whereas footshock-induced velocity was normal during a secondary non-instrumental shock-sensitivity test (**Extended Data Fig. 2f,g**). These data show that when a preferred option is associated with risk of punishment, mice adjust their choice to a less valuable option in a manner requiring neuronal activity in BLA.

**FIGURE 2.**
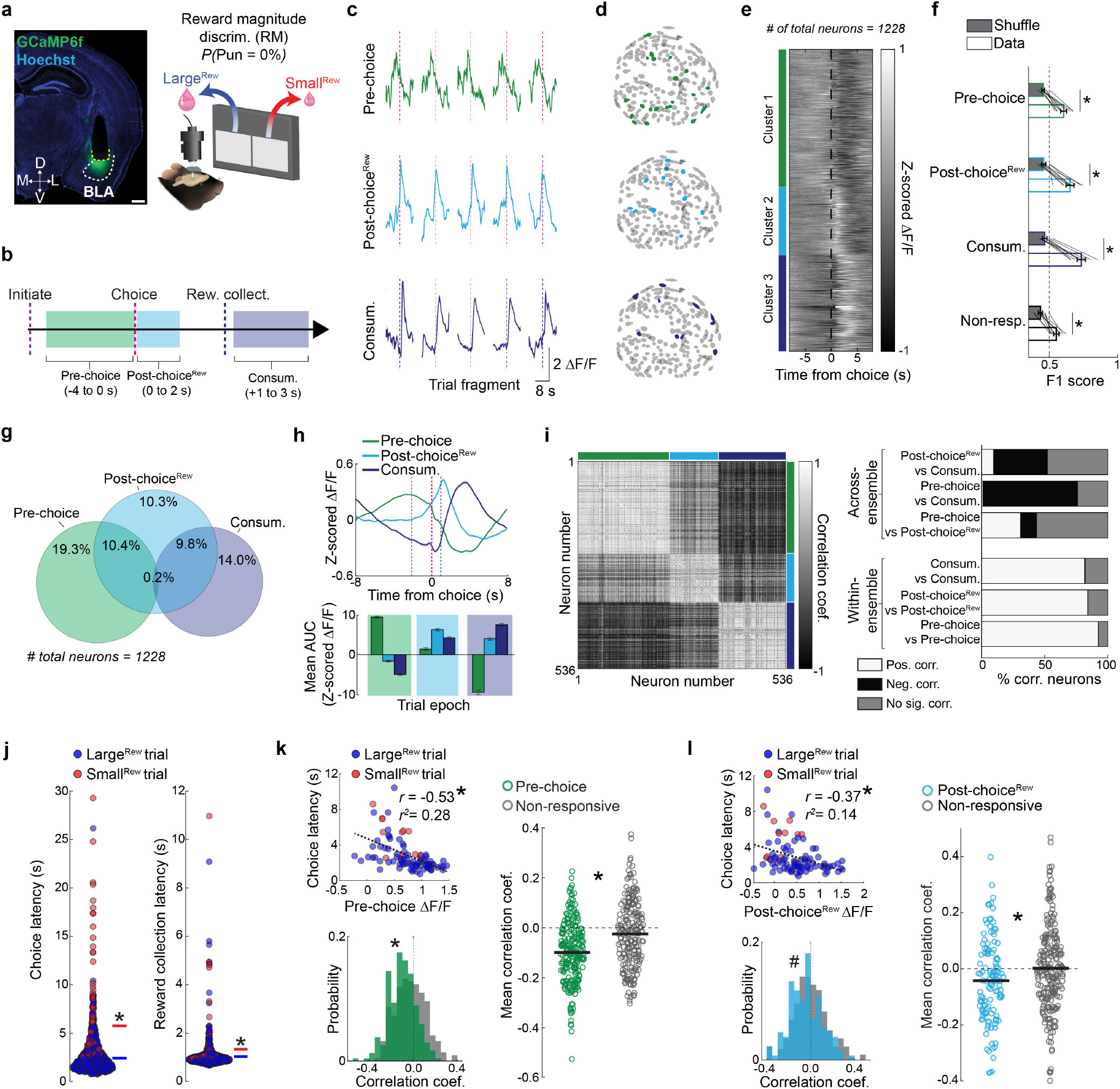
Dedicated but opposing BLA neuronal ensembles encoding choice and consumption. (**a**) Example GRIN lens location targeting BLA (left) (Scale bar: 500 µm); cartoon depicting in vivo microendoscope Ca^2+^ imaging during reward magnitude discrimination (RM) testing (right). (**b**) Epochs defining neuronal-encoding of Pre-choice, Post-choice^Rew^, and Consum. events. (**c**) Event-aligned activity over five trials for a representative cell from each ensemble. (**d**) Topography of ensemble neurons in the lens field of view from the same example mouse. (**e**) Heatmap displaying choice-aligned activity from all imaged neurons (n=1228 neurons), grouped by k-means cluster (colored bars indicate ensemble identity). (**f**) F1 scores from decoding. For all ensembles (and non-responsive neurons), empirical data from each ensemble-specific epoch (non-responsive epoch was set to Consum. window) could be decoded from shuffle (Shuffle vs Data for each ensemble: all p<0.05, paired-samples t-tests). (**g**) Degree of overlap between Pre-choice, Post-choice^Rew^ and Consum. ensembles. (**h**) Ensemble activity aligned to choice, with median trial initiation time (purple dashed lines), time of choice (pink dashed line), and the median time of reward collection (dark blue dashed line) indicated (top); epoch ensemble AUCs (bottom). (**i**) Pair-wise Pearson correlation matrix organized by ensemble identity (colored bars indicate ensemble identity) (left). Summary of correlations across and within-ensemble (right). (**j**) Individual trial data (n = 900 events) indicating choice (left) and reward collection (right) latency, with shorter mean latencies for Large^Rew^ versus Small^Rew^ (both p<0.001, unpaired t-tests). (**k**) Negative correlation between choice latency and Pre-choice ensemble activity in a representative neuron (circles represent single-trial data; r=-0.53, r2=0.28, p<0.001, Pearson correlation) (upper left). Negatively-skewed distribution of binned correlation coefficients for Pre-choice (green), relative to non-responsive (grey), neurons (k(452)=0.24, p<0.0001, Kolmogorov-Smirnov test) (lower left). Higher overall negative correlation for Pre-choice versus non-responsive neurons (t(452)=6.21, p<0.001, unpaired t-test) (right). (**l**) Analysis as in **k**, for Post-choice^Rew^ neuron activity (representative negatively-correlated neuron, r=-0.37, r2=0.14, p<0.05, Pearson correlation) (upper left). The correlation distribution of Post-choice^Rew^ neurons (light blue) tended to be negatively-skewed versus non-responsive (grey) neurons (k(344)=0.14, p=0.0665, Kolmogorov-Smirnov test) (lower left). Higher negative correlation for Post-choice^Rew^, relative to non-responsive, neurons (t(344)=2.43, p<0.02, unpaired t-test) (lower right). n=10 mice for all comparisons. For full reporting of statistics, see **Supplementary Table 1**. Data mean ± SEM. *p<0.05, # p<0.07. Reward water drop (**a**, doi.org/10.5281/zenodo.3925935) graphic adapted from scidraw.io.

### Distinct performance-related choice and consumption ensembles

We next examined the native choice-related activity of BLA neurons by virally expressing the Ca^2+^ indicator GCaMP6f in BLA neurons and conducting 1-photon cellular-resolution Ca^2+^ imaging via a chronically implanted gradient refractive index (GRIN) lens (**Fig. 2a, left, Extended Data Fig. 3a**). To characterize task-related BLA activity, we inspected the peri-choice activity of 1228 neurons (extracted using constrained non-negative matrix factorization for microendoscopic data, CNMF-E; Zhou et al. 2018) recorded from 10 mice during Late RM (well-trained mice, no punishment risk) (**Fig. 2a, right, Extended Data Fig. 3b,c**). Using the event-sequence inherent to the task’s trial-structure, we isolated neuronal dynamics in three epochs corresponding to the Pre-choice, Post-choice^Rew^ and reward consumption (Consum.) periods (**Fig. 2b**).

We allocated active neurons to one or more epoch-related ensemble (approach as in Jimenez et al. 2018) (**Fig. 2c**), which had no clear spatial organization within our imaging fields of view (Fig. 2d). Unbiased K-means clustering demarcated neurons into comparable ensembles with activity tiling the entire peri-choice period (**Fig. 2e, Extended Data Fig. 3d**). We found that ensemble characteristics held irrespective of choice-type, in that the dynamics of each ensemble were broadly similar when separately analyzing Large^Rew^ and Small^Rew^ trials after equating the number of trials for each of the two choice-types (**Extended Data Fig. 3e**). Moreover, linear classifiers trained on the neuronal data from each epoch had higher decoding performance for ensemble neurons than shuffled data (**Fig. 2f**). On examining the relationship between ensembles, however, we found that Pre-choice and Consum. ensembles were almost completely non-overlapping, whereas Post-choice^Rew^ neurons partially overlapped with the other two (**Fig. 2g**). Pre-choice and Consum. encoding neurons were also oppositely modulated, such that Consum. ensemble activity was low when Pre-choice activity was high, and vice versa (**Fig. 2h**). Likewise, these two ensembles showed highly anti-correlated activity patterns (**Fig. 2i**).

We next sought to relate the activity of ensemble neurons to behavioral performance by leveraging trial-wise variations in choice and reward-collection latency (**Fig. 2j**). Interestingly, we found that the activity of Pre-choice and Post-choice^Rew^, but not Consum., neurons correlated with faster choices (**Fig. 2k,l, Extended Data Fig. 3f**). Consum. neuron activity was also unrelated to reward consumption duration (**Extended Data Fig. 3g**). Of note, we found no clear behavioral evidence that mice became sated or fatigued across the experimental session (**Extended Data Fig. 3h-k**), implying that the correlations between choice-related activity and latencies were not driven by these factors. Hence, these data suggest that neuronal activity around the time of choice may reflect the incentive value of the preferred outcome and the vigor associated with choosing that option. Importantly, similar results were obtained when reproducing these analyses using neuronal Ca^2+^ data recorded during the safe RDT trial-block (**Extended Data Fig. 3l-n**).

In sum, these initial imaging data show that, under safe conditions, dedicated and distinct BLA neuronal ensembles encode behaviorally relevant aspects of choice for a desired outcome.

### Punishment encoding shapes willingness to choose

We next performed Ca^2+^ imaging during the RDT to ask how BLA neurons encoded punishment and whether this activity reflected riskrelated behavioral change (**Fig. 3a**). Receipt of the footshock punisher caused the activation of a substantial portion of neurons (Post-choice^Pun^; **Fig. 3b**), many of which were re-activated on a second RDT conducted days later (**Extended Data Fig. 4a**), suggesting that these neurons stably represented punishment. Post-choice^Pun^ active neurons were differentiable from those responsive to reward, as demonstrated by opposing activity profiles and largely non-overlapping ensemble membership from Consum. neurons identified either during the RDT (**Fig. 3c**) or during a risk-free touchscreen-based progressive ratio test (**Extended Data Fig. 4b**). At the behavioral level, we found that the risk-related slowing of task-related behaviors was positively correlated with Post-choice^Pun^ neuronal activity. Specifically, activity of this ensemble was higher on trials when reward collection was slower (**Fig. 3e,f**) but unrelated to another choice-related index of behavioral slowing, subsequent trial initiation latency (**Extended Data Fig. 4c,d**). By contrast, the negative correlation between Pre-choice neuronal activity and choice latency that was evident on safe trials (see **Fig. 2k, Extended Data Fig. 3f**) was absent under risk (**Extended Data Fig. 4e**).

**FIGURE 3.**
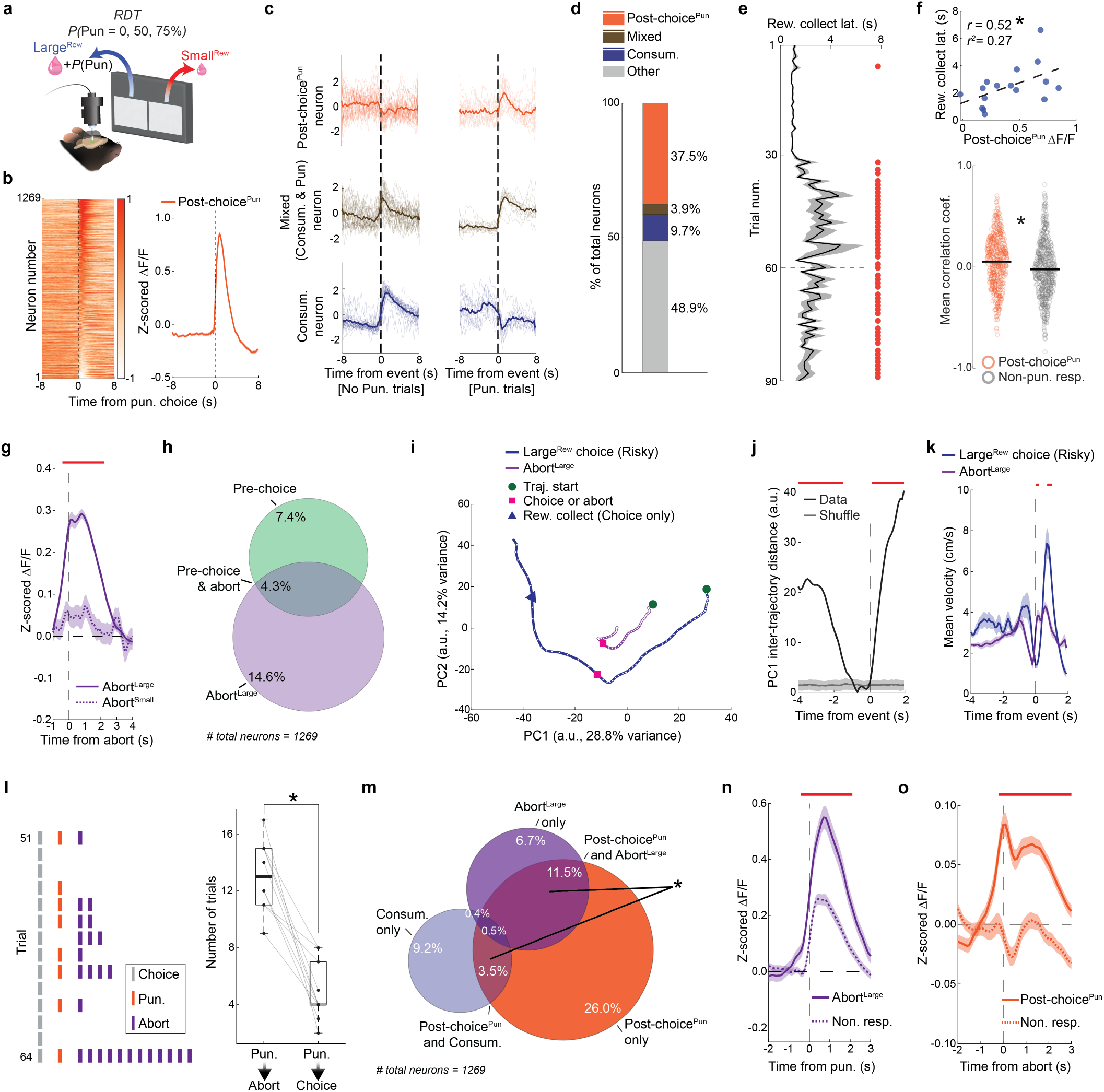
BLA ensembles encode a punisher and its effect on choice. (**a**) Cartoon depicting in vivo microendoscope Ca^2+^ imaging of BLA neurons during RDT testing. (**b**) Choice-aligned heatmap of neuronal activity on punished trials (sorted by peak activity 0-2 s post-choice) (left) and corresponding mean Post-choice^Pun^ ensemble activity (right). (**c**) Trial mean (thick line) and individual trial (thin lines) activity from three representative neurons aligned to choice on non-punished (left) and punished (right) Large^Rew^ trials. (**d**) Proportion of neurons encoding Post-choice^Pun^, Mixed (responsive on punished and non-punished trials), and Consum. only. (**e**) Longer reward-collection latency on trials during risky trial blocks (horizontal dashed lines demarcate blocks, red circles indicate trials where p<0.001, bootstrapped confidence interval computed from mean choice latencies in the safe block). (**f**) Positive correlation between reward collection latency and Post-choice^Pun^ activity (circles represent single-trial data; r=0.52, r2=0.27, p<0.05, Pearson correlation) (upper). Higher positive correlation for Post-choice^Pun^ than non-responsive, neurons (t(1093)=-4.85, p<0.001, unpaired t-test) (lower). (**g**) Higher activity during aborts directed at the Large^Rew^ (Abort^Large^), not Small^Rew^ (Abort^Small^), response-window during risky blocks. (**h**) Low overlap between Abort^Large^ and Pre-choice (i.e., trial executed) ensembles during risky blocks. (**i**) Divergent event-aligned PCA trajectories for Abort^Large^ and Large^Rew^ neuronal activity during risky blocks (excluding punished trials). (**j**) Euclidean distances of PC1 between empirical (black line) and shuffled (grey line) data. (**k**). Similar mean locomotor velocity prior to Abort^Large^ and Large^Rew^ choice. Significant velocity differences only emerge after time 0s due to velocity spiking prior to reward collection following Large^Rew^ choice, but not Abort^Large^. (**l**) Rasterized trials selected from a representative RDT session illustrating that choices (grey lines) that are punished (orange lines) tend to be followed by at least one abort (purple lines) (left). More trials in which punishment was followed by an abort(s) versus a choice (t(9)=7.17, p<0.001, paired t-test) (right). (**m**) Greater overlap between Post-choice^Pun^ and Abort^Large^ neurons than between Post-choice^Pun^ and Consum. neurons (*χ*^2^(1)=49.03, p<0.0001, Chi-square test). (**n**) Higher activity of Abort^Large^, but not abort non-responsive, neurons during punishment delivery. (**o**) Elevated activity of Post-choice^Pun^ ensemble, but not punishment non-responsive, neurons during the peri-abort period. Red horizonal lines in **g,j,n**, and **o** indicate samples that significantly differ between groups, p<0.001, permutation test). For full reporting of statistics, see **Supplementary Table 1**. Data mean ± SEM. *p<0.05. Reward water drop (**a**, doi.org/10.5281/zenodo.3925935) graphic adapted from scidraw.io.

These data suggest that the representation of the punished outcome in BLA neurons persists beyond the immediate aversive sensation of shock to influence subsequent behavior. To explore this possibility, we asked whether BLA activity was related to a major emergent feature of behavior under risk – aborted choices (see **Fig. 1h,l**). We found that a subset of BLA neurons were active when mice exhibited aborts directed at the Large^Rew^ option (Abort^Large^), but not on the rare occasions when aborts were directed at the Small^Rew^ option (**Fig. 3g**). Interestingly, Abort^Large^ neurons were largely distinct from neurons active prior to choices that were fully executed (i.e., Pre-choice neurons) (**Fig. 3h**). Differential encoding of aborted and executed choice was further evidenced by decomposing mean activity traces into orthogonal principal components, which revealed divergent neuronal trajectories for the two actions, despite each being performed at a similar speed up to their occurrence (**Fig. 3i-k**).

Given that abort responses seem to be driven by a process distinct from that which drives choice, we next examined the relationship between punishment experience and the propensity to abort. Behaviorally, we found that punished trials were more likely to be followed by at least one abort (**Fig. 3l**). At the neural level, abort encoding neurons were more likely to be responsive to punishment than to reward – i.e., abort neurons overlapped more with the Post-choice^Pun^ than Consum. ensemble (**Fig. 3m**) – and encoded punishment more strongly than did neurons non-responsive to aborts (**Fig. 3n**). Post-choice^Pun^ encoding neurons were also more strongly reactivated at the time when choice was aborted, as compared to neurons that were inactive during punishment (**Fig. 3o**).

Together, these observations suggest that the aversive properties of the punished outcome directly influence aborted actions and are neurally represented in BLA at the time mice abort their choice for the Large^Rew^ option, shaping the willingness to choose a desired but risky option.

### Risk revises representation of choice

Next, we turned to assessing how punishment affected the ensemble representations of choice in BLA that were observed during safety (see **Fig. 2**). By comparing ensemble-specific neuronal activity across the safe (‘Safe-identified’) and risky (‘Risky-identified’) RDT trial-blocks, we found that risk broadly reduced the activity of Safe-identified Pre-choice, Post-choice^Rew^ and Consum. ensembles (**Fig. 4a,b**). The loss of Safe-identified ensemble activity was not due to there being relatively more Small^Rew^ choices during risky blocks as this effect was still evident when limiting our analysis to only Large^Rew^ trials (**Extended Data Fig. 5a**).

**FIGURE 4.**
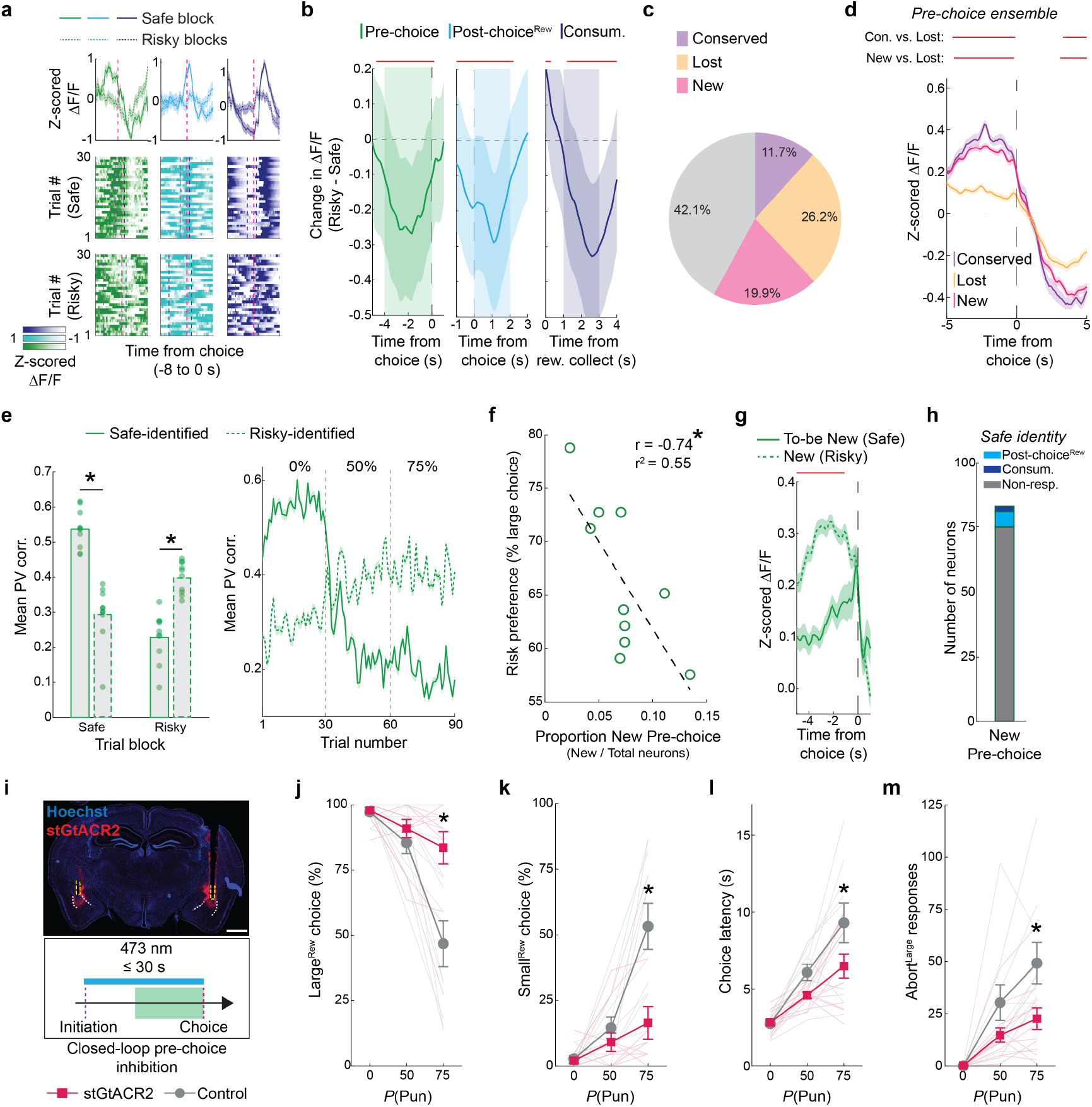
Risk of punishment transforms BLA neuronal representation of choice. (**a**) Choice-aligned activity from representative Pre-choice (green), Post-choice^Rew^ (light blue), and Consum. (dark blue) neurons during safe (solid line) and risky (dashed line) blocks (upper). Choice-aligned heatmaps showing loss of ensemble-specific encoding during risky (lower), relative to safe (center), blocks. (**b**) Decreased neural activity in Safe-identified ensembles during Risky blocks, indicated by a decrease in difference score (Risky – Safe) (red lines indicate samples significantly different from 0, p<0.001, bootstrapped confidence interval). (**c**) Proportion of neurons exhibiting Conserved, Lost, or New ensemble identity across safe and risky RDT blocks. (**d**) Mean activity traces for Pre-choice ensemble neurons that were Conserved (maintained identity across Safe to Risky blocks), Lost (lost identity across safe to risky blocks), or New (exhibited identity on risky not safe blocks) (red horizonal line indicates samples different between groups indicated at left, p<0.001, permutation tests). (**e**) PV correlations were lost for Safe-identified (solid green border) neurons on risky blocks (t(9)=12.94, p<0.001, paired t-test), and emerged for Risky-identified (dashed green border) neurons on risky blocks (t(9)=-4.69, p<0.001, paired t-test) (left). Mean trial-wise PV correlations for Safe and Risky-identified neurons (right). (**f**) Negative correlation between risk preference and the proportion of New Pre-choice neurons during risky blocks. (**g**) New Pre-choice neurons were pre-choice quiescent on safe blocks (To-be New) (red horizonal line indicates samples different between groups, p<0.001, permutation test). (**h**) Most New Pre-choice neurons were non-responsive on safe trials. (**i**) Representative stGtACR2 expression and optic fiber tip locations in BLA (dashed yellow line; scale bar: 1 mm) (upper). Schematic of closed-loop optogenetic (blue line) manipulation during the pre-choice period (green shading) (lower). (**j,k**) Increased Large^Rew^ (**j**), and decreased Small^Rew^ (**k**), choice during risky, relative to safe, trials-blocks following stGtACR2-mediated silencing (stGtACR2 vs Control: both p<0.001 on 75% risky block; ANOVA interaction: F(2,42)=10.73, p<0.001). (**l,m**) stGtACR2-mediated BLA silencing led to shorter risky-block choice latency (stGtACR2 vs Control: p<0.05 on 75% risky block; ANOVA interaction: F(2,42)=3.37, p<0.05) (**l**) and fewer Abort^Large^ responses (stGtACR2 vs Control: p<0.05 on 75% risky block; ANOVA interaction: F(2,42)=4.30, p<0.05). Multiple comparison tests for j-m were Tukey’s post-hoc tests conducted after significant Treatment (stGtACR2 vs Control) x Trial Block (0, 50, 75%) (**m**) ANOVA interaction. For full reporting of statistics, see **Supplementary Table 1**. Data mean ± SEM. *p<0.05.

Interestingly, these risk-related reductions in activity masked more complex changes occurring in subpopulations of BLA neurons. Specifically, we found that whereas ∼26% of Safe-identified neurons lost (Lost) their event-related activity on risky trial-blocks, approximately half that number (∼12%) retained it (Conserved) and ∼20% of neurons showed newly acquired event encoding (New) (**Fig. 4c**), with similar changes evident for Pre-choice, Post-choice^Rew^ and Consum. ensembles (**Extended Data Fig. 5b**). Comparing the profile of choice-aligned activity in these subpopulations on risky trials showed that New and Conserved neurons were similar to one another, but, as expected, different (i.e., more active) from Lost neurons (**Fig. 4d, Extended Data. Fig. 5c**). Lastly, there was less conservation and greater loss of ensemble identity on across the RDT session than the Late RM session (**Extended Data Fig. 5d**).

**FIGURE 5.**
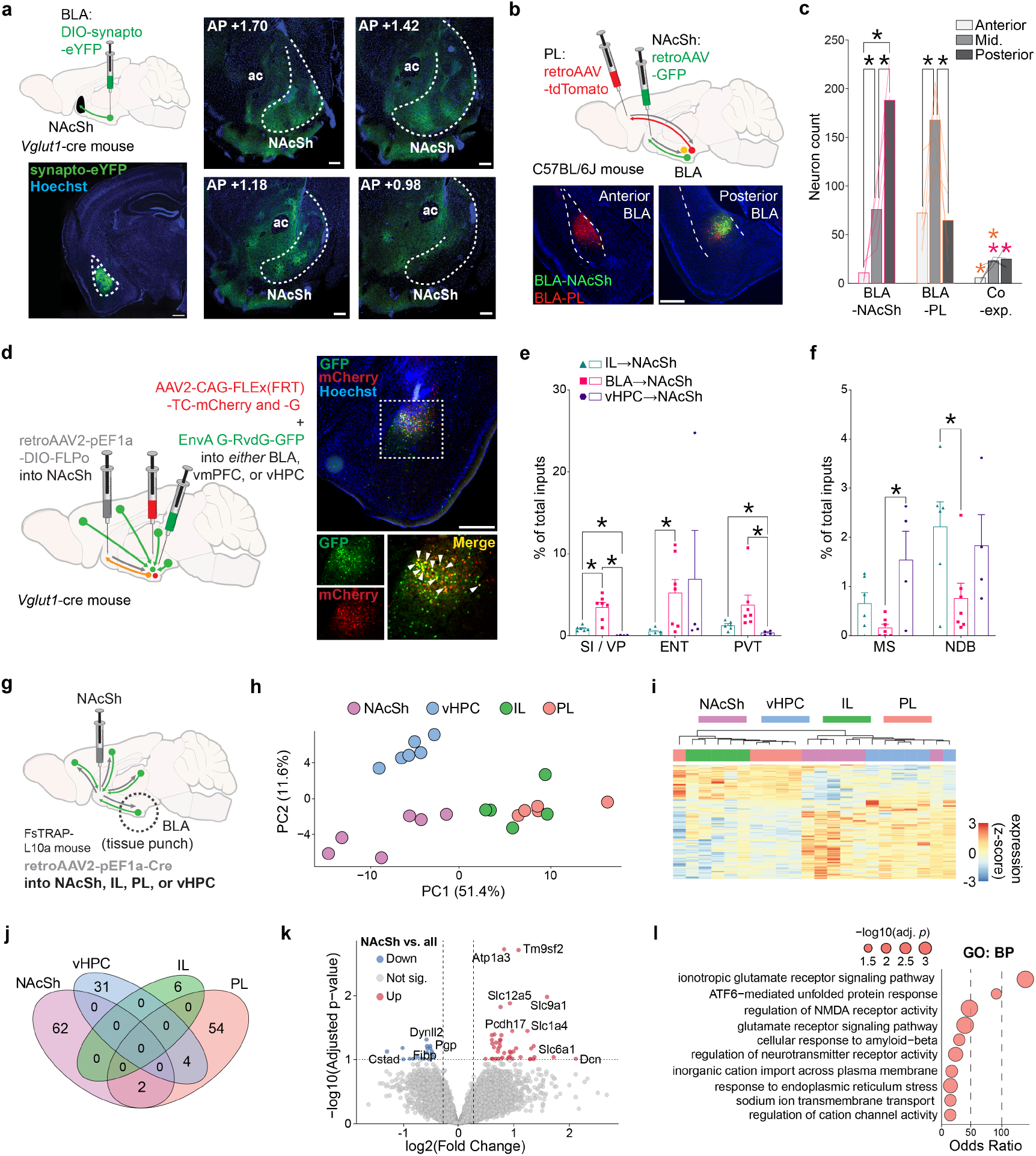
Mapping outputs and inputs and transcriptional profile of BLA neurons projecting to nucleus accumbens shell. (**a**) BLA-originating excitatory (Vglut1-cre) synaptic terminals labeled in NAcSh. Scale bars: 500 µm (BLA image), 200 µm (NAcSh images). (**b**) Schematic of retrograde viral tracing approach for labeling NAcSh- and PL-projecting BLA neurons, with representative images of virus expression. Scale bars: 500 µm. (**c**) NAcSh-projecting BLA neurons originated in mostly posterior BLA whereas PL-projecting neurons tended to be localized mid-way (Mid) between the anterior and posterior limits of BLA. Minimal co-expression (Co-exp) of NAcSh- and PL-projecting BLA neurons. (**d**) Schematic of rabies viral tracing approach for labeling inputs to BLA^→^NAcSh neurons. Scale bar: 500 µm. (**e**) Denser input from ventral pallidum (VP), entorhinal cortex (ENT), and the paraventricular thalamus (PVT) to BLA^→^NAcSh versus vHPC^→^NAcSh (VP: t(6.01)=6.15, p<0.001; PVT: t(6.11)=2.86, p<0.05, unpaired t-tests) or vmPFC^→^NAcSh neurons (VP: t(6.64)=4.54, p<0.005; ENT: t(6.13)=6.13, p<0.05, unpaired t-tests). (**f**) Lesser input from medial septum (MS) and the nucleus of the diagonal band (NDB) to BLA^→^NAcSh than vHPC^→^NAcSh (MS: t(4.20)=3.14, p<0.05, unpaired t-test) or vmPFC^→^NAcSh neurons (NDB: t(8.29)=2.45, p<0.05, unpaired t-test). (**g**) Schematic for TRAPseq tagging of EGFP to the large ribosomal subunit (L10) in specific BLA output populations. (**h**) Principal Component Analysis (PCA) of differentially expressed genes (DEGs) identified across BLA projection targets using a likelihood ratio test (LRT, FDR < 0.05) shows clear separation by projection group. (**i**) Heatmap of z-scored expression values for DEGs confirms distinct transcriptional profiles across projection-defined groups. (**j**) Venn diagram illustrating overlap of DEGs identified in one-vs-all DESeq2 contrasts. The majority of DEGs are specific to a single projection pathway. (**k**) Volcano plot highlighting top DEGs from the NAcSh versus all comparison. (**l**) Gene Ontology enrichment analysis showing the top 10 enriched terms. Point size indicates –log10(adjusted p-value); the x-axis shows the odds ratio. Data mean ± SEM. *p<0.05. Analyses in e and f were unpaired t-tests with Welch correction. For full reporting of statistics, see **Supplementary Table 1**. Hamilton Syringe (**a, b, d, g**, doi.org/10.5281/zenodo.7679042) and Mouse Brain (**a, b, d, g**, doi.org/10.5281/zenodo.3925909) graphics adapted from scidraw.io.

To assess the degree to which ensemble activity was affected by the introduction of risk at the population level, we constructed population vectors (PVs) reflecting the mean activity for neurons within each ensemble-specific epoch identified during safe or risky trial-blocks. PVs were then correlated with session-long Ca^2+^ activity and the mean correlation during each epoch-specific sub-window was calculated for every trial. We found that trial-wise PV correlations for Safe-identified Pre-choice ensembles were strong in the safe block but declined during risky blocks, suggesting a loss in pre-choice encoding across the population (**Fig. 4e, left**). This diminished encoding was balanced by an increase in the strength of encoding in the Risky-identified Pre-choice ensemble that emerged during the transition from the safe to risky block (**Fig. 4e, right**). Comparable changes were observed for Post-choice^Rew^ and Consum. ensembles, indicative of a widespread shift in population-level dynamics across blocks (**Extended Data Fig. 5e,f**).

These findings suggest that ensemble reorganization and population-remapping related to the dramatic change in choice behavior that occurs with the introduction of risk. However, it remained unclear which facet of these changes related most strongly to behavioral change. Strikingly, we found that the size of the New Pre-choice ensemble was significantly negatively correlated with the level of risk preference on an individual mouse level, such that risk-averse mice gained more New Pre-choice neurons on risky trial-blocks, whereas risky mice gained fewer (**Fig. 4f**). This relationship was specific to New Pre-choice neurons and was not apparent for either Lost or Conserved neurons or for any subset of the Post-choice^Rew^ and Consum. ensembles (**Extended Data Fig. 5g**). By retrospectively tracking neurons from the risky to safe trial-block, we were able to show that ‘To-be New’ Pre-choice cells were largely unresponsive on safe trials and were mainly derived from a pool of non-responsive neurons (**Fig. 4g,h**).

These imaging data imply that dynamic changes in the Pre-choice BLA neuronal representation causally contribute to risk-related alterations in behavior. To test this hypothesis, we expressed within BLA neurons the high efficiency inhibitory opsin, stGtACR2 (Mahn et al. 2018), or a control fluorophore, and bilaterally implanted optic fibers to deliver blue light (5 mW, 473 nm) to BLA during the pre-choice period of each RDT trial on risky and safe trial-blocks (**Fig. 4i, Extended Data Fig. 5h**). We found that stGtACR2-expressing mice made more and faster Large^Rew^ choices and were less likely to perform aborted Large^Rew^ actions during risky trial-blocks, as compared to controls (**Fig. 4j-m**). Both groups displayed similar behavioral responses to the footshock punisher within the RDT session and during a subsequent shock-sensitivity test (**Extended Data Fig. 5i,j**). These results are consistent with a key contribution of Pre-choice BLA neuronal encoding to risk-driven shifts in RDT performance.

These convergent imaging and optogenetic data are consistent with the importance of BLA representational dynamics as a key neural substrate of risk-driven adaptations in behavior.

### Anatomical-genetic profile of NAcSh-projecting BLA neurons

We next turned to the question of how BLA representations of punished choice are read out downstream, focusing on a BLA to medial nucleus accumbens shell (NAcSh) pathway previously linked to both reward-seeking and aversive states (Ambroggi et al. 2008, Beyeler et al. 2016 2018, Kim et al. 2016, Britt et al. 2012, Folkes et al. 2020, Lind et al. 2023, Millan et al. 2017, Namburi et al. 2015, Shen et al. 2019, Stuber et al. 2011, Zhang et al. 2021). Using viral labeling, we found that glutamatergic (Vglut1-expressing) BLA projection neurons densely innervated the anterior and mid-anterior extent of the medial NAcSh, as revealed by the presence of synaptophysin-eYFP expressing synaptic terminals arriving from BLA (**Fig. 5a**). This pattern of BLA^→^NAcSh innervation was corroborated by comparison to Allen Connectivity Data (Oh et al. 2014) using Brain Street View (https://github.com/Julie-Fabre/brain_street_view; **Extended Data Fig. 6a**). Using retroAAVs to express retrogradely-trafficked fluorophores into two distinct BLA projection targets (**Fig. 5b, Extended Data. Fig. 6b**), we observed that NAcSh-projecting neurons were mostly localized in the posterior BLA and were largely distinct from neurons that projected to the prelimbic prefrontal cortex (PL; **Fig. 5c**), in line with prior reports (Brog et al. 1993, Groenewegen et al. 1999, Huang et al. 2021, Kita and Kitai 1990, McGarry and Carter 2017, Shinonaga et al. 1994).

**FIGURE 6.**
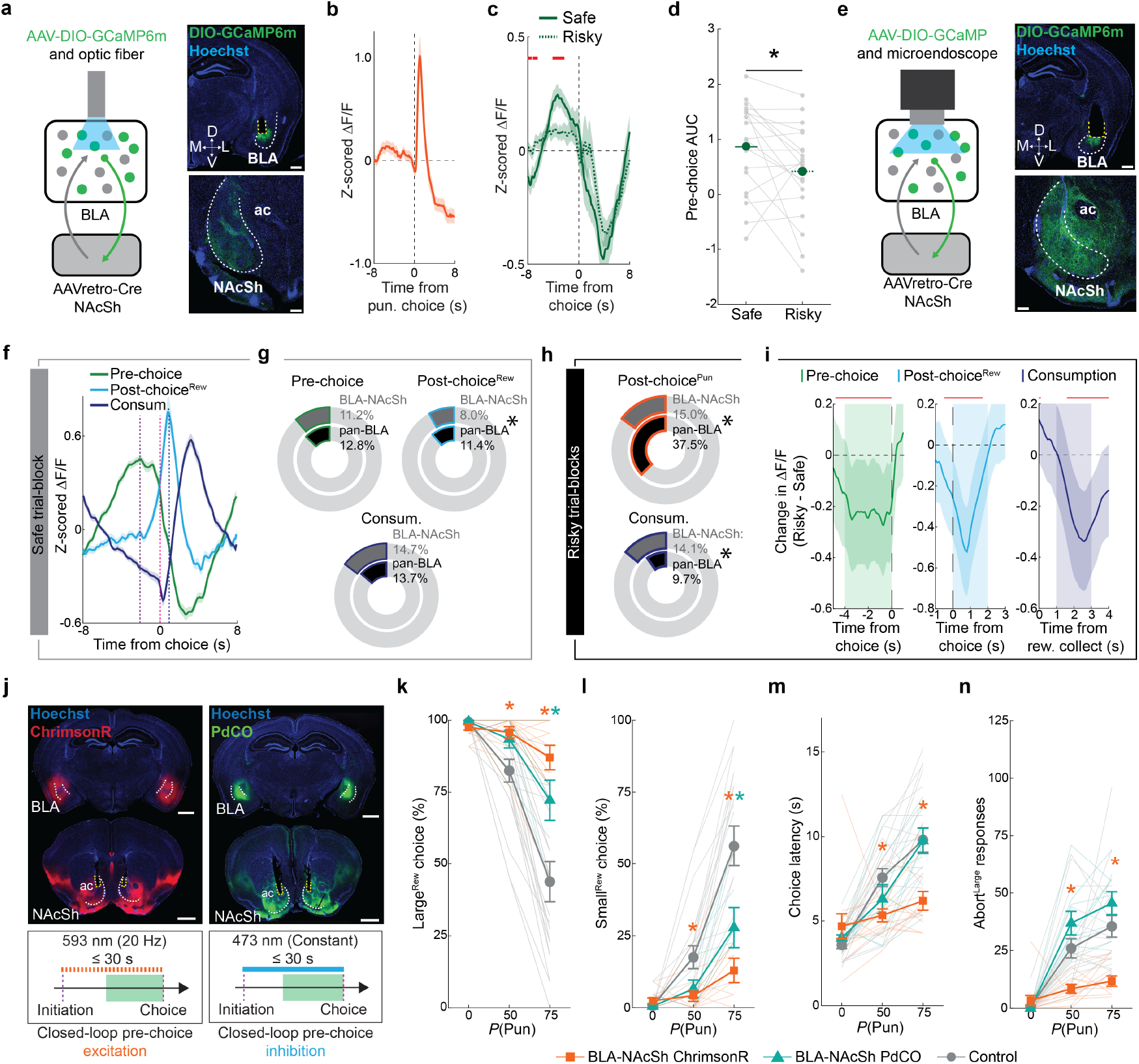
BLA encoding of punished choice is readout to nucleus accumbens shell (NAcSh). (**a**) Cartoon depiction of in vivo fiber photometry in NAcSh-projecting BLA neurons (left). DIO-GCaMP expression and optic fiber tip (dashed yellow line) in BLA (top right). GCaMP expression in NAcSh. Scale bars: 500 µm (BLA), 200 µm (NAcSh) (bottom right). (**b**) BLA^→^NAcSh activity during punishment. (**c**) Choice-aligned activity during safe and risky blocks (red horizonal line indicates samples different between Safe and Risky blocks, p<0.001, permutation tests). (**d**) Lesser trial-averaged pre-choice activity (AUC) on risky versus safe trial blocks (t(19)=2.26, p<0.05, paired t-test). (**e**) Cartoon depiction of in vivo microendoscope Ca^2+^ imaging in NAcSh-projecting BLA neurons (left), with example of GCaMP expression in BLA (top right) and NAcSh (bottom right) Scale bars: 500 µm (BLA), 200 µm (NAcSh). (**f**) Choice-aligned activity of event-defined ensemble neurons, with median trial initiation time (purple dashed line), time of choice (pink dashed line), and the median time of reward collection (dark blue dashed line) indicated (comparable to pan-BLA imaging data show in **Fig. 2h**). (**g**) Proportionally fewer safe-block-identified Post-choice^Rew^ BLA^→^NAcSh neurons versus pan-BLA imaging (z=2.55, p<0.05, two-proportion z-test). (**h**) Proportionally fewer Post-choice^Pun^ (z=12.85, p<0.0001, two-proportion z-test), but more Consum. neurons (z=2.77, p<0.006; two-proportion z-test) in BLA^→^NAcSh than pan-BLA imaged neurons on risky blocks. (**i**) Decreased (safe-block-identified) ensemble activity during risky blocks (red lines indicate samples significantly different from 0, p<0.001, bootstrapped confidence interval). (**j**) Representative images of opsin expression in BLA and NAcSh and optic fiber tip locations (dashed yellow lines) in BLA. Scale bars: 1 mm. (top). Schematic of closed-loop optogenetic manipulation during the pre-choice period (green shading) using ChrimsonR (bottom left, orange dashed line indicates illumination) or PdCO (right, blue line indicates illumination) (bottom right). (**k,l**) Optogenetic manipulation of BLA^→^NAcSh produced risk-insensitive patterns of choice (ANOVA interaction: F(4,78)=10.48, p<0.0001) as evidenced by increased Large^Rew^ (**k**) and decreased Small^Rew^ (**l**) choice during risky, relative to safe, blocks following ChrimsonR-mediated activation (ChrimsonR vs Control: p<0.05 on both risky blocks) or PdCO-mediated inhibition (PdCO vs Control: p<0.05 on 75% risky block). (**m, n**) Shorter choice latency (ANOVA interaction: F(4,78)=7.87, p<0.001) (**m**) and fewer Abort^Large^ responses (ANOVA interaction: F(4,78)=7.97, p<0.001) (**n**) on both risky blocks in the ChrimsonR group (ChrimsonR vs Control: all p<0.05 on both risky blocks) but not PdCO group. Data mean ± SEM. Multiple comparison tests for **k-n** were Dunnett’s tests comparing the Control group to the ChrimsonR or PdCO groups, conducted following significant Treatment (Control, ChrimsonR, PdCO) x Trial Block (0, 50, 75%) ANOVAs. For **k-n**, color of * indicates significant difference (p<0.05) between specific opsin condition and Control. For full reporting of statistics, see **Supplementary Table 1**. *p<0.05.

We next examined whether the BLA^→^NAcSh pathway is embedded within distinct circuits as compared to other NAcSh-projecting pathways. To do so, we mapped monosynaptic inputs to glutamatergic (Vglut1-expressing) BLA^→^NAcSh neurons using a viral-genetic rabies tracing approach (Schwarz et al. 2015) (**Fig. 5d, Extended Data Fig. 6c,d**). Rabies-positive BLA^→^NAcSh-projecting cells were enriched in the paraventricular nucleus of the thalamus (PVT), entorhinal cortex (ENT), and the ventral pallidum (VP) relative to NAcSh-projecting neurons in infralimbic (IL) prefrontal cortex or ventral hippocampal (vHPC) (**Fig. 5e, Extended Data Fig. 6d**). Conversely, rabies-labeled inputs from the medial septum (MS) and nucleus of the diagonal band (NDB) were sparser in the BLA^→^NAcSh-projecting population (**Fig. 5f**). Other multi-synaptic circuits were identified that differed across NAcSh input pathways, with control experiments verifying that rabies-labeled inputs were appropriately projection- and genetically-defined (**Extended Data Fig. 6c-j**).

To provide insight into the molecular characteristics of BLA^→^NAcSh neurons, we transcriptionally profiled these neurons, compared against a separate set of NAcSh-projecting regions, via translating ribosome affinity purification (Ekstrand et al. 2014) (**Fig. 5g, Extended Data Fig. 7a**). This approach identified 131 differentially expressed genes (DEGs) across all pathways, based on group-wise comparisons using the likelihood ratio test (LRT) (**Fig. 5h,i**). To identify projection-specific markers, we performed pairwise comparisons using Wald’s test, which revealed 62 DEGs enriched in BLA^→^NAcSh neurons but absent in BLA^→^PL, BLA^→^IL, and BLA^→^vHPC populations (**Fig. 5j**). These genes included Dcn, Nrcam, Slc1a4, and Slc9a1 (**Fig. 5k, Supplementary Table 2**). Gene ontology analysis highlighted functional enrichment for glutamatergic synaptic transmission, and KEGG pathway analysis revealed significant associations with nicotine and amphetamine addiction, cAMP signaling, and glutamatergic transmission (**Fig. 5i, Extended Data Fig. 7c, Supplementary Table 3**).

To further refine projection-specific gene signatures, we applied sparse Partial Least Squares Discriminant Analysis (sPLS-DA) in a one-vs-rest classification framework. This supervised, multivariate approach selects a minimal subset of genes that best distinguish each projection group, with top-ranked sPLS-DA features across regions largely reproducing the projection-based clustering observed with PCA (**Extended Data Fig. 7d**). By comparing multivariate markers with univariate DEGs, we found that a substantial proportion of sPLS-DA features overlapped with DESeq2-derived DEGs, with the greatest concordance observed in the BLA^→^PL and BLA^→^vHPC groups (**Extended Data Fig. 7e,f**). sPLS-DA-identified genes included Slc9a1 (NAcSh), Ifrd1 (PL), Sst (vHPC), and 4930524B15Rik (IL) (**Extended Data Fig. 7g**).

Together, these data characterize the distinct anatomical profile of BLA^→^NAcSh neurons and suggest these neurons may utilize distinct molecular machinery to regulate synaptic function.

### NAcSh-projecting BLA neurons mediate adaptation to risk

After characterizing NAcSh-projecting BLA neurons, we assessed whether neurons in this pathway functionally contributed to RDT performance. First, by employing in vivo fiber photometry to measure bulk changes in neuronal Ca^2+^ (**Fig. 6a, Extended Data Fig. 8a-c**), we found that BLA-NAcSh projecting neurons were responsive to post-choice punishment and that their activity during the pre-choice period of the risky blocks was depressed relative to the safe RDT block (**Fig. 6b**). Importantly, this latter effect was not seen across blocks in which risk was absent (i.e., Late RM) (**Extended Data Fig. 8d-f**). These activity patterns indicate that BLA^→^NAcSh neurons are responsive to risk-related variables in a manner consistent with our pan-BLA imaging results, although our ability to resolve activity during distinct epochs (e.g., post-choice and consumption) was curtailed by the measurement of bulk (single pixel) fluorescence measurement.

To extend these photometry data, we injected a retroAAV-Cre in NAcSh and a Cre-dependent GCaMP6m in BLA before implanting a GRIN lens above BLA to conduct in vivo cellular-resolution Ca^2+^ imaging of BLA^→^NAcSh neurons during the RDT (**Fig. 6e, Extended Data Fig. 9a-f**). Analyzing the activity of 653 neurons from 20 mice revealed that, during the safe trial-block, sub-populations of BLA^→^NAcSh neurons could - as in our pan-BLA imaging dataset - be characterized as being active during the pre-choice, post-choice and consumption periods (**Fig. 6f**). The size of each ensemble was also similar to those imaged pan-neuronally, although BLA^→^NAcSh was slightly less enriched in Post-choice^Rew^ neurons (**Fig. 6**). The three ensembles also displayed many of the same characteristics during safe blocks that we observed in our pan-BLA imaging experiment (see **Fig. 2 and Extended Data Fig. 3l-n**), including ensemble opposition, inter-ensemble correlation, and an inverse relationship between choice latency and peri-choice activity (**Extended Data Fig. 9g-j**).

When examining data from the risky blocks, differences were evident in the encoding properties of the BLA^→^NAcSh neurons as compared to the entire BLA population. Specifically, we found proportionally fewer Post-choice^Pun^ neurons in the BLA^→^NAcSh population, as compared to the BLA population overall (**Fig. 6h, top**), and a greater proportion of Consum. neurons (**Fig. 6h, bottom**) – together suggesting that the BLA^→^NAcSh pathway may be biased towards positive valence encoding. Aside from these differences in ensemble size, how-ever, BLA^→^NAcSh neurons generally displayed risk-related changes in activity similar to those we observed during pan-BLA neuronal imaging, including event-related activity during the pre-choice, post-choice and consumption epochs on safe trials that was significantly blunted on risky trials (**Fig. 6i**).

Given these imaging data demonstrate that BLA^→^NAcSh neurons undergo dynamic changes in response to risk, we employed optogenetics to causally interrogate the functional contribution of these dynamics to RDT performance. To this end, we expressed either the high efficiency excitatory opsin ChrimsonR (Klapoetke et al. 2014), the synaptic transmission-suppressing G-protein-coupled opsin PdCO (Wietek et al. 2024), or an opsin-negative fluorophore control in BLA neurons and bilaterally implanted optic fibers to deliver red (ChrimsonR) or blue (PdCO) light at NAcSh specifically during the pre-choice period of each RDT trial (**Fig. 6j, Extended Data Fig. 10a,b**). Interestingly, we found that mice expressing either opsin made more Large^Rew^ choices than controls on the riskiest RDT block (**Fig. 6k,l**). Additionally, ChrimsonR-mediated activation (but not PdCO-mediated inhibition) of the BLA^→^NAcSh pathway during the pre-choice period attenuated risk-related increases in both choice latency (**Fig. 6m**) and the performance of Abort^Large^ responses (**Fig. 6n**). Lastly, on a touchscreen PR task, ChrimsonR-mediated activation dampened, rather than increased, motivation, excluding the possibility that the increased riskiness produced by this manipulation was due to enhanced motivation for the reward (**Extended Data Fig. 10c**). Thus, our observation that bi-directional manipulation of BLA^→^NAcSh impairs RDT performance suggests that the perturbation of normal BLA^→^NAcSh activity dynamics is sufficient to prevent choice flexibility.

In sum, these photometry and imaging data indicate that the NAcSh is a key output through which the BLA mediates the ability of animals to adapt behavior under risk of punishment.

## 3 DISCUSSION

Here we present evidence implicating BLA in reward-guided choice in the absence and presence of punishment risk. By first establishing how BLA ensembles respond during safe reward-seeking, we provide support for a recently-outlined framework suggesting the opposite modulation of BLA ‘action’ and ‘consumption’ neurons in self-paced operant behavior (Courtin et al. 2022). This ensemble opposition adds to growing evidence that a general characteristic of amygdala function is the existence of distinct populations that encode mutually exclusive behavioral states, which, as we show here, is also maintained under risk of punishment. These data are also broadly consistent with the purported contribution of BLA to energizing performance in an outcome-specific manner on other tasks (Ambroggi et al. 2008, Ghods-Sharifi et al. 2009, Piantadosi et al. 2017, Servonnet et al. 2020, Tye and Janak 2007). This function was evident in the absence of risk (i.e., on safe RDT trials) in the form of a relationship between choice-related BLA neuronal activity (specifically in Pre-choice and Post-choice^Rew^ ensembles) and the vigor with which animals made choices. However, given BLA inhibition in the present study was without behavioral effect on safe trials, the BLA appears to be dispensable for maintaining actions in the pursuit of a high-value outcome when animals are well trained (Balleine et al. 2003, Ghods-Sharifi et al. 2009, Pelloux et al. 2013, Piantadosi et al. 2017).

Our results add additional support for the consistent finding that rodents with lesions or pharmacological inactivation of BLA maintain reward-seeking despite punishment (Choi and Kim 2010, Ishikawa et al. 2020, Jean-Richard-Dit-Bressel and McNally 2015, Marchant et al. 2019, Orsini et al. 2015, Pelloux et al. 2013, Piantadosi et al. 2017, Verharen et al. 2019): here, chemogenetic or optogenetic inhibition of BLA neurons increased riskiness by maintaining preference for a high-value reward and preventing the production of aborted choices. These findings broadly align with the known contribution of BLA to regulating actions based on expected outcomes and modifying these actions when outcomes change (Balleine and Killcross 2006, Courtin et al. 2022, Janak and Tye 2015, Kyriazi et al. 2018, Salzman and Fusi 2010, Sharpe and Schoenbaum 2016, Shen et al. 2019). Notably, the increased risky behavior produced by chemogenetic inhibition of BLA was associated with a blunted response to the footshock in the RDT, raising the possibility that reduced sensitivity to the punisher contributed to the increased riskiness. However, individual differences in riskiness did not relate to shock-sensitivity (i.e., **Extended Data Fig. 1f**), suggesting that this explanation is unlikely. In addition, when footshock sensitivity was measured independent of reward-seeking, neither chemogenetic nor optogenetic inhibition of BLA had any impact on responsivity. This dissociation aligns with recent conceptual accounts proposing that punishment insensitivity in humans occurs due to an inability to causally link actions with punished consequences and alter behavior accordingly, despite normal sensitivity to the aversiveness of the punisher (Jean-Richard-dit Bressel et al. 2021, Jean-Richard-Dit-Bressel et al. 2023). Thus, our findings support the idea that BLA may be an important neural substrate underlying the capacity to connect actions with punishment.

In this context, prior studies have shown that BLA representations of stimuli/responses become similar to an associated aversive outcome when animals repeatedly experience them together (Grewe et al. 2017, Gründemann et al. 2019, Zhang et al. 2021, Zhang and Li 2018), including during punishment, wherein neuronal responses (measured using photometry) to punishment become comparable to those occurring during the punishment-producing action (Jean-Richard-dit Bressel et al. 2022). This convergence of encoding has led to the hypothesis that BLA representation of the aversive properties of an action’s punished outcome may underlie reductions in the execution of that action, in a similar manner to that shown to occur in NAc-projecting mPFC neurons (Kim et al. 2017) and D2-expressing neurons within NAc (Zalocusky et al. 2016).

We provide several lines of evidence supporting this model in BLA. First, we observed that mice were more likely to abort than execute a choice on the trial following a punished outcome and, furthermore, that separable BLA ensembles encoded aborted and executed choices: a dissociation reminiscent of the differential BLA single-unit encoding of conditioned approach versus avoidance responses (Kyriazi et al. 2018) and completed versus aborted foraging actions in the presence of a pseudopredator (Amir et al. 2015). Second, we demonstrate that punishment-responsive BLA neurons are active during aborted choices and that neurons that are responsive to punishment preferentially encode aborted over executed choices. Taken together, these findings suggest that by representing an aversive experience encountered around the time of choice, BLA neuronal activity modifies an animal’s behavioral strategy in a manner that mitigates the potential for future punishment.

Another key finding here is that these representations of an aversive outcome emerge quickly in BLA. We found that when animals encountered punishment, BLA neurons rapidly (within a single test session) modified how choice-related information was represented. This was evident when comparing neuronal activity on safe versus risky trial-blocks at both the single neuron (as losses and gains in ensemble membership) and population level (as a change in population vector). These findings extend recent evidence that BLA neuronal encoding in mice and monkeys rapidly remaps when instrumental action-outcome expectations are altered by contingency degradation or outcome revaluation/devaluation (Courtin et al. 2022, Fustiñana et al. 2021, Gründemann et al. 2019, Morrison and Salzman 2010).

Comparing the present findings with earlier work, it is noteworthy that, despite manipulating outcomes in dissimilar ways (punishment versus reinforcer devaluation), both the current and a prior study in mice (Courtin et al. 2022) found that changes in the expected outcome resulted in the emergence of new action/choice-encoding neurons. These convergent findings suggest that the incorporation of newly encoding neurons might be a common feature of BLA representational alterations occurring with violation of an expected outcome value (as has also been seen in regions such as orbitofrontal cortex, Schoenbaum et al. 1998 2000). Moreover, we went on to show that the relative size of a specific population of neurons - those that acquired pre-choice responsivity during the risky trial-blocks - was predictive of the degree to which mice became risky versus risk-averse. Hence, the dynamic integration of newly-encoding neurons into the pre-choice BLA representation could be an important neural adaptation contributing to the risk-driven shift away from the preferred choice option.

In support of this idea, optogenetically inhibiting BLA neurons during the pre-choice period caused mice to maintain choice for the high-value reward despite the risk of punishment. Though this experiment could not isolate the contribution of new Pre-choice neurons to risk aversion per se, the results are in line with the importance of pre-choice neuronal dynamics to risk-driven shifts in actions. In contrast, an earlier study conducted in rats found that pre-choice optogenetic BLA inhibition decreased, rather than increased, riskiness (Orsini et al. 2017). This apparent discrepancy likely reflects substantial differences in the psychological constructs under investigation in the two studies, as mediated by task experience: here, we studied mice learning about risk over a single RDT experience, whereas earlier work examined rats retrieving a well-learned behavioral strategy (Orsini and Simon 2020). Our data therefore provide new insight into the acquisition and immediate updating of actions based on a novel experience of punishment. The fact that there are marked experience-dependent differences in the effect of BLA inhibition implies a hidden complexity to the region’s role in risky decision-making which awaits further investigation.

In part, this complexity may reflect engagement of different BLA output circuits. We obtained evidence that BLA neurons projecting to the NAcSh are at least one of the major functional output populations underpinning risk-related behavioral flexibility. By leveraging open-source data and in-house viral tracing, we confirmed that BLA neurons target medial NAcSh, are located in posterior BLA, and are separate from PL-projectors, largely supporting prior work (Brog et al. 1993, Groenewegen et al. 1999, Huang et al. 2021, Kita and Kitai 1990, McGarry and Carter 2017, Shinonaga et al. 1994). Fiber photometry and cellular-resolution microendoscopy revealed task-related activation (e.g., during punishment and reward-consumption) and risk-related alterations (e.g., pre-choice decreases) in BLA^→^NAcSh neuronal activity, while altering the normal activity of this pathway, by either optogenetically exciting or silencing BLA^→^NAcSh neurons, promoted riskiness. These effects are in keeping with data from other loss-of-function studies that implicate BLA^→^NAcSh neurons in risk-taking (Bercovici et al. 2018, Truckenbrod et al. 2023, van Holstein et al. 2020) and the suppression of alcohol-seeking after extinction, cue omission or reward unavailability (Millan et al. 2015 2017). There are, however, other studies showing that chemogenetic or optogenetic activation of BLA^→^NAcSh neurons is reinforcing and drives reward-seeking actions (Britt et al. 2012, Folkes et al. 2020, Lafferty et al. 2020, Stuber et al. 2011). One potential explanation to reconcile these opposite observations is that the ‘risky’ behavior induced here by BLA^→^NAcSh activation stems from enhanced motivation for the large reward. However, this interpretation is unlikely given that ChrimsonR-mediated BLA^→^NAcSh activation here mildly diminished objective measures of motivation measured on a PR task (**Extended Data Fig. 10c**). Hence, the role of this pathway in regulating rewarded actions appears to be complex. Nonetheless, the contribution of BLA^→^NAcSh neurons to adapting behavior in response to risk suggests this population may be dysfunctional in situations characterized by excessive or deficient sensitivity to risk.

Relatedly, our sequencing results are in keeping with prior investigations suggesting that BLA neurons that project to distinct regions are also partially genetically distinct (Kim et al. 2016, Namburi et al. 2015, O’Leary et al. 2020, Zhang et al. 2021). This characterization of the genetic profile of BLA^→^NAcSh neurons provides a basis for potentially identifying therapeutic intervention targets to mitigate neuropsychiatric abnormalities in risk sensitivity. In addition, we report that BLA^→^NAcSh neurons receive unique inputs as compared to other accumbens afferents, including from PVT. PVT and its direct projection to NAcSh have been shown to control aspects of reward-seeking during threat (Choi et al. 2019, Choi and McNally 2017, Engelke et al. 2021), implying that there may be parallel polysynaptic pathways available for the regulation of such behavior.

In sum, we have shown that when faced with the risk of punishment, mice strategically alter their behavior to reduce the likelihood of experiencing an aversive outcome while still obtaining a lower value reward. We find that these behavioral adaptations are associated with rapid, dynamic changes in choice-related BLA neuronal representations and show that the incorporation of newly pre-choice encoding neurons predicts the degree to which behavior adapts in response to risk. Collectively, these data provide novel evidence for a key contribution of BLA to modifying behavioral actions when an expected rewarded outcome is associated with potential risk of punishment. Given amygdala abnormalities are evident in humans exhibiting impaired risk-related decision-making (Bechara et al. 1999, Brand et al. 2007, Weller et al. 2007), these findings could have implications for understanding the neural factors underlying variation in risk for neuropsychiatric disorders characterized by excesses in risk taking or aversion.

## 4 METHODS

### Subjects

All experiments were conducted in compliance with the National Institutes of Health Guide for the Care and Use of Laboratory Animals and approved by the local National Institute on Alcohol Abuse and Alcoholism Animal Care and Use Committee. Subjects were male C57BL/6J (strain #00664) mice purchased from The Jackson Laboratories (Bar Harbor, ME, USA), male Vglut1-Cre mice (strain #023527) purchased from The Jackson Laboratories and bred in-house to female wild-type C57BL/6J mice to produce heterozygous mice for experimentation, and male homozygous Rosa26^fsTRAP^ mice (strain #022367) purchased from The Jackson Laboratories and bred in-house to female C57BL/6J mice to produce heterozygous mice for experimentation. Vglut1-Cre mice were used to selectively target BLA excitatory neurons (see details below); Vglut1 is strongly expressed in excitatory projection neurons (Fremeau et al. 2001, Fujiyama et al. 2001, Lein et al. 2007, Zeisel et al. 2018). Rosa26^fsTRAP^ mice were used to transcriptionally profile distinct BLA neurons projecting to distinct outputs (see details below Rosa26^fsTRAP^ mice enable Cre-dependent expression of GFP on the large ribosomal subunit, Heiman et al. 2014 2008). Mice were group-housed (maximum 4 mice/cage) in a temperature and humidity-controlled vivarium (12/12 h light/dark cycle, lights on at 6 AM) and allowed ad libitum access to food and water, unless specified below. Mice were between 2-6 months of age at the time of experimentation.

### Interface implantation and virus infusion

#### General surgical procedures

Mice were anesthetized with vaporized isoflurane (induction at 5%, maintenance at 1-3%) and secured into a stereotaxic frame (Kopf Instruments, Tujunga, CA, USA). Body temperature was maintained at 37°C using a self-regulating electrical heater. The head was shaved and cleaned with betadine followed by alcohol before the skin was incised or trimmed with scissors. Craniotomies were made with 0.7 or 0.9-mm drill bits attached to a handheld drill (Ideal Micro-Drill). Infusions were conducted over 10 minutes using a 32 ga Hamilton syringe (Neuros Syringe, Model 7000.5, 0.5 µL, Hamilton Company, Reno, NV, USA) attached to a manual injector (Model 1772 Universal Holder, Kopf Instruments) or a syringe pump (UMP3 with Micro4 pump controller, World Precision Instruments, Sarasota, FL, USA). Following injection, the syringe was left in place for a further 10 minutes to ensure fluid infusion was complete.

For surgeries involving chronic optical interface implantation, a pilot hole was drilled at the posterior portion of the incision to secure a single stainless-steel skull screw (00-90 x 1/16 inch, Model AMS90/1P, Antrin Miniature Specialties, Fallbrook, CA, USA). Following surgery, mice were given 5 mg/kg of the analgesic Ketoprofen in Lactated Ringers Solution and singly-housed to ensure healing of the surgical site and integrity of the implant. All animals were monitored daily and allowed to recover for a minimum of 1 week prior to experimentation.

#### In vivo 1-photon imaging

To image calcium (Ca^2+^) activity in BLA neurons, an AAVdj-hSynGCaMP6f (∼700 nl, titer 1.0 x 10^13^, Stanford Gene Vector and Virus Core, Stanford, CA, USA) was unilaterally infused into the left hemisphere BLA. To image Ca^2+^ activity specifically in nucleus accumbens shell (NAcSh)-projecting BLA neurons, an rAAV2-retro-EF1α-Cre (∼300 nl, titer 8.35 x 10^12^, Salk Institute Viral Vector Core, La Jolla, CA, USA) was unilaterally infused into the NAcSh (coordinates from bregma/skull AP: +1.55, ML: 0.62, DV: -4.61) and a Cre-dependent AAVdj-EF1α-DIO-GCaMP6m (∼700 nl, titer 4.0 x 10^12^, University of North Carolina Viral Vector Core, Chapel Hill, NC, USA) or AAV5-syn-FLEX-jGCaMP8f-WPRE (∼700 nl, titer 1.97 x 10^13^, Addgene, Watertown, MA, USA) was unilaterally infused into BLA (coordinates from bregma/skull, AP: -1.50; ML: +/- 3.23; DV: -5.00).

For both imaging experiments, a gradient refractive index (GRIN) lens (0.6 mm diameter, 7.3 mm length, Inscopix Inc. part #1050-004413, or 0.66 mm diameter, 7.5 mm length, Inscopix Inc. part #1050-005422, Inscopix, Mountain View, CA, USA) with a pre-attached baseplate was implanted unilaterally above the BLA (coordinates from bregma/skull, AP:-1.52; ML: +/- 3.27; DV: -4.90). To affix the lens and baseplate to the skull, a small amount of super glue (KrazyGlue) was applied to the skull and was reinforced with 1-2 layers of Metabond (Parkell, Edgewood, NJ, USA) followed by black dental acrylic (Coralite Dental Products, Skokie, IL, USA) to minimize light penetrance. The pan-BLA imaging experiment used n=10 mice, and the BLA^→^NAcSh imaging experiment used n=20 mice.

#### In vivo fiber photometry

To measure bulk Ca^2+^ activity in NAcSh-projecting BLA neurons using fiber photometry, an AAV containing retrogradely traveling rAAV2-retro-EF1α-Cre (∼300 nl, titer 8.35 x 10^12^, Salk Institute Viral Vector Core) was unilaterally infused into NAcSh (coordinates from bregma/skull AP: +1.55, ML: 0.62, DV: -4.61) and a Cre-dependent AAVdj-EF1α-DIO-GCaMP6m (∼300 nl, titer 4 x 10^12^, University of North Carolina Viral Vector Core) was unilaterally infused into BLA (coordinates as above). A fiber-optic implant (400 µm, 0.57 numerical aperture, B280-4419-5, Doric Lenses, Quebec, QC, CA) was lowered into place above the BLA (unilaterally, with the implantation hemisphere counter-balanced across mice, coordinates from bregma/skull, AP: -1.55; ML: +/- 3.24; DV: -4.90) and affixed using a thin layer of KrazyGlue glue followed by application of layers of pink and then black dental acrylic. This experiment used n=20 mice.

#### In vivo chemogenetics

To chemogenetically inhibit BLA neurons, an AAV5-hSyn-hM4Di-mCherry (∼300 nl, titer 2.5 x 10^13^; Addgene) or an AAV5-hSyn-mCherry control (∼300 nl, titer 1.7 x 10^13^, Stanford Gene Vector and Virus Core) was bilaterally infused into BLA (coordinates from bregma/skull, AP: -1.50; ML: +/- 3.24; DV: -5.00). This experiment used n=15 hM4Di-expresing mice and n=14 mCherry-expressing mice (1 of these mice could not be used for the saline-control RDT session, resulting in n=13 for that session).

#### In vivo optogenetics

To optogenetically inhibit BLA cell bodies, AAV1-CAMKIIα-stGtACR2.0-FusionRed (Mahn et al. 2018) (∼200 nl, titer 7 x 10^12^, Addgene) or an EYFP control (∼200 nl, titer 3.6 x 10^12^, University of North Carolina Viral Vector Core) was bilaterally infused (coordinates from bregma/skull, AP: -1.50; ML: +/- 3.24; DV: -5.00). Optic fibers (cut to 5 mm length, 200 µm diameter, 0.39 numerical aperture; Thorlabs, Newton, NJ, USA) were bilaterally implanted targeting BLA (coordinates from bregma/skull, AP: -1.55; ML: +/- 3.25; DV: -4.75). This experiment used n=13 stGtACR2-expressing mice and n=10 EYFP-expressing mice.

To optogenetically excite or inhibit BLA axons in NAcSh, an AAV containing either the excitatory opsin ChrimsonR (Klapoetke et al. 2014) (AAV5-syn-ChrimsonR-tdT, ∼200 nl, titer 8.2 x 10^12^, Addgene), the inhibitory opsin PdCO (Wietek et al. 2024) (AAV5-hSyn-PdCO-EGFP-WPRE, ∼200 nl, titer 1.4 x 10^13^, Addgene) or an mCherry control (AAV5-hSyn-mCherry, ∼200 nl, titer 2.3 x 10^13^, Addgene) was bilaterally infused into BLA (coordinates from bregma/skull, AP: -1.50; ML: +/- 3.24; DV: -5.00). Optic fibers (cut to 5 mm length, 200 µm diameter, 0.39 numerical aperture; Thorlabs) were bilaterally implanted targeting NAcSh (coordinates from bregma/skull, AP: +1.55; ML: +/- 1.34; DV: -4.10, at an angle of 10°). This experiment used n=13 ChrimsonR-expressing mice, n=9 PdCO-expressing mice, and n=20 mCherry-expressing mice.

#### Anterograde labeling of BLA terminals in NAcSh

To visualize BLA synaptic terminals in the NAcSh, Vglut1-Cre+ mice were unilaterally infused with an AAV containing an eYFP-tagged Cre-dependent version of the anterograde synaptic label, synaptophysin (∼200 nl, AAV8.2-hEF1α-DIO-synaptophysin-eYFP, titer 2.1 x 10^13^, Gene Delivery Technology Core, Massachusetts General Brigham, Boston, MA, USA) into BLA (coordinates from bregma/skull, AP: -1.50; ML: +/- 3.24; DV: -5.00). Mice were perfused 1 month after injection. Images from a single representative mouse are displayed after verifying expression within NAcSh was consistent across n=4 mice (data not shown).

#### Retrograde labeling of BLA outputs to NAcSh and PL

To quantify the degree of overlap between BLA outputs to NAcSh and a comparison region, the prelimbic (PL) cortex, and to describe their anterior-posterior expression level, a retrogradely-expressed AAV containing either GFP (∼300 nl, AAVrg-CAG-GFP, titer 7 x 10^12^, Addgene) or tdTomato (∼300 nl, AAVrg-CAG-tdTomato, titer 2.1 x 10^13^, Addgene) was unilaterally infused into either NAcSh (coordinates from bregma/skull, AP: -1.55; ML: +/- 1.32; DV: -4.53, at an angle of 10°) or PL (coordinates from bregma/skull AP: +1.95, ML: +/-1.00, DV: -1.90, at an angle of 20°). This experiment used n=4 mice.

#### Transsynaptic rabies tracing

To identify upstream regions monosynaptically projecting to BLA^→^NAcSh, IL^→^NAcSh, or vHPC^→^NAcSh neurons, the ‘cell-type specific tracing the relationship between input and output’ (cTRIO) method was employed (Schwarz et al. 2015). To target excitatory output neurons within the BLA, Vglut1-Cre+ mice were unilaterally infused with a retrogradely-expressed retroAAV2-pEF1α-DIO-FLPo-WPREhGHpA (∼500 nl, titer 6.2 x 10^13^ ; Addgene) into NAcSh (coordinates from bregma/skull AP: +1.55, ML: 0.62, DV: -4.61). Additionally, a 1:1 mixture of Flp-dependent AAVs (∼500 nl total) containing the EnvA receptor TVA fused to mCherry (AAV2-CAG-FLEx(FRT)-TC, ∼250 nl, original titer 1.6 x 10^13^; Addgene, packaged by ViGene, Rockville, MD, USA) and the rabies glycoprotein (AAV2-CAG-FLEx(FRT)-G, ∼250 nl, original titer 1.4 x 10^13^; Addgene, packaged by ViGene) was unilaterally infused into BLA (coordinates from bregma/skull, AP: -1.50; ML: +/- 3.23; DV: -5.00), IL (coordinates from bregma/skull AP: +1.90; ML: +/- 1.40; DV: -2.90, at an angle of 20°), or vHPC (coordinates from bregma/skull AP: -3.10; ML: +/- 3.22; DV: -4.53). Approximately 5 weeks later, an EnvA-pseudotyped G-deleted version of the rabies virus (Chatterjee et al. 2018) (EnvA G-deleted Rabies-GFP, ∼500 nl, titer 5.45 x 10^8^; Salk Institute Viral Vector Core) was infused into BLA, IL, or vHPC (coordinates as above), and mice were perfused 5 days later. This experiment used mice n=7 BLA^→^NAcSh mice, n=6 IL^→^NAcSh mice, and n=4 vHPC^→^NAcSh mice.

#### Surgical procedures for TRAPseq transcriptional profiling

To transcriptionally profile BLA neuronal outputs, heterozygous Rosa26^fsTRAP^ mice were bilateral infused with rAAV2-retro-EF1α-Cre (∼350 nl, titer 2.1 x 10^13^, Addgene) into NAcSh (coordinates from bregma/skull AP: +1.55, ML: +/-0.62, DV: -4.61), PL (coordinates from bregma/skull AP: +1.95, ML: +/-1.00, DV: -1.90, at an angle of 20°), IL (coordinates from bregma/skull AP: +1.90; ML: +/- 1.40; DV: -2.90, at an angle of 20°), or vHPC (coordinates from bregma/skull AP: -3.10; ML: +/- 3.22; DV: -4.53) and perfused 1 month later.

### Touchscreen testing

#### Apparatus

Mice were tested using the Bussey-Saksida Touch Screen System (Model 80614; Lafayette Instruments, Lafayette, IN, USA). Each chamber featured a trapezoidal arena with a reward port and house light on one side and a touchscreen monitor on the other (**Fig. 1a**). Touches on the monitor were restricted (by a black plexiglass mask) to two response-windows (7 x 7.62 cm) within which the stimuli (1x 6.5 cm^2^ white square per window) were displayed. The arena floor was a stainless-steel grid which was connected to a scrambled shock generator (Campden Instruments, Loughborough, Leicestershire, UK). A computer running Whisker Server and ABET II Touch software was used to control the inputs and outputs for each chamber. Behavioral programs (described below) were custom written within ABET II Touch.

#### Pretraining

Following recovery from surgery, mice were gradually (over the course of 1 week) restricted to 80-85% of their free feeding weight. Mice first underwent daily sessions (1 session per day) on a 5-phase pre-training protocol to develop familiarity with the touchscreen manipulanda, as previously described (Glover et al. 2020). In phase 1, mice were acclimated to milkshake reward (∼1 ml, Nesquik Strawberry Milkshake, Vevey, Switzerland), delivered into a plastic weigh boat in their home cage. In phase 2, mice were trained during a 60-minute test session in the touchscreen chamber to associate a 2-second, 65 dB pure tone (hereafter, reward tone) with reward (∼5 µl) delivered into the illuminated reward port. Mice collecting 30 rewards within 30 minutes progressed to phase 3. During phase 3, mice were required to initiate a trial by breaking an infrared beam located in the illuminated reward port, which caused the reward port light to extinguish and a visual stimulus (randomly selected from a stimulus library) to be displayed on 1 of the 2 touchscreen windows. Touching the stimulus or waiting 20 seconds resulted in illumination of the reward port, presentation of the reward tone, and reward delivery. Mice progressed to phase 4 by collecting 30 rewards in 30 minutes (regardless of how many screen touches were made). Phase 4 was identical to phase 3, except that mice were required to touch the visual stimulus to receive reward. The criterion to advance was the same as for phase 3. Phase 5 was identical to phase 4, except that touching the blank touchscreen response-window now produced a 20-second ‘timeout’ period during which the house light was extinguished and mice were unable to initiate a new trial. Mice advanced from this phase when they correctly touched the response-window containing the stimulus on >75% of 30 trials and completed the session within 30 minutes.

#### Reward magnitude discrimination (RM)

After pre-training, mice began daily RM discrimination sessions. Sessions were organized into 3 discrete 30-trial blocks. A block began with 8 forced-choice trials: on 4 of the trials, the visual stimulus (6.5 cm^2^ white square) was presented on the left screen; on the other 4 trials, the same stimulus was presented on the right screen, according to a pseudorandom programmed sequence. During the next 22 free-choice trials, the same stimuli were presented on both screens simultaneously (**Fig. 1a**). For each mouse, either the left-presented or right-presented stimulus was designated as the large reward (∼20 µl), with the other stimulus designated as the small reward (∼5 µl). For each trial-type, stimulus presentation occurred when the mouse initiated a trial by breaking the infrared beam located in the reward port.

A trial was completed when the mouse made a touchscreen response at a stimulus, resulting in reward delivery, along with illumination of the reward port and delivery of the reward tone (**Supplemental Video 1**). If no response was made within 30 seconds, the stimuli were extinguished and the trial was scored as an omission. Omitted trials did not count towards the 8 forced and 22 free choice trials within each block – following an omission, the trial entered the 8 second inter-trial interval (ITI; during which new trials could not be initiated) state upon which the trial was repeated upon initiation. On completed trials, collection of the reward broke the reward port infrared beam which extinguished the reward port light and initiated the 8 second ITI. Following the ITI, the reward port light was illuminated to signal that a new trial could be initiated. Mice were given daily test sessions until reaching a performance criterion of responding at the large reward stimulus on >90% of free-choice trials and completing the session in <2000 seconds. To ensure discrimination performance was robust and efficient, mice were trained for a minimum of 8 sessions.

#### Risky decision making test (RDT)

After reaching RM criterion, mice underwent RDT testing. The RDT was structured identically to the reward magnitude discrimination sessions, consisting of 3 blocks of 30 trials (8 forced-choice, and 22 free-choice trials per block) except that the selection of the large reward option could produce (in addition reward delivery) a 0.1 mA scrambled foot-shock at a probability of 0% (block 1), 50% (block 2) or 75% (block 3) (**Supplemental Video S2**). To allow mice to experience footshock at these probabilities, forced-choice trials in each block delivered shocks at the exact probability associated with each trial block (0%: 0 shocks, 50%: 2 shocks out of 4 total large reward trials, 75%: 3 shocks out of 4 total large reward trials). During free-choice trials for each trial block, shocked trials were selected from a probability distribution centered around 0, 50, or 75%. Selection of the small reward was never associated with footshock.

#### Touchscreen procedures for fiber photometry and microendoscopic imaging

On each test session from the start of RM discrimination training, mice were connected to a patch cable or miniaturized microscope, connected to a commutator to allow for unimpeded movement (for further details, see Piantadosi et al. 2025). Fiber photometry data were collected during each RM discrimination session. To minimize photobleaching, 1-photon imaging data was collected only during the first (‘Early RM’) and last (‘Late RM’) RM session. Photometry and imaging were conducted over two RDT sessions, each separated by a minimum of 2 RM discrimination re-training sessions to reestablish performance criterion.

#### Touchscreen procedure for chemogenetics

On attaining RM discrimination criterion, mice received a habituation i.p. saline injection 30 minutes prior to 1 or more additional RM sessions until criterion was re-attained. RDT was conducted on the next session, 30 minutes after i.p. injection of clozapine N-oxide (CNO; 3 mg/kg, BioTechne Corporation, Minneapolis, MN, USA). Mice were then retrained daily (drug-free) to RM criterion (median number of sessions: 9, range: 7-24 sessions), before receiving a second RDT test conducted 30 minutes after i.p. saline injection.

#### Touchscreen procedure for optogenetics

On each test session from the start of RM discrimination training, mice were connected to bilateral patch cables that were connected to a commutator to allow for unimpeded movement (for further details, see Piantadosi et al. 2025). Once mice achieved the RM discrimination criterion, they underwent an optogenetic manipulation session where pre-choice activity was manipulated. For stGtACR2 (**Fig. 4i-m**) or PdCO (**Fig. 6j-n**) mice (or their respective controls), illumination (∼5 mW) of a solid-state 473 nm wavelength laser (MBL-III-473, Opto Engine LLC, Midvale, UT, USA) was timed to occur upon trial initiation on free-choice trials only. For ChrimsonR (or their respective control) mice (**Fig. 6j-n**), illumination (∼2 mW) of a solid-state 593.5 nm wavelength laser (MGL-F-593.5, Opto Engine LLC) was timed as above, but pulsed at 20 Hz (5 ms pulse duration). Laser illumination was terminated by a choice or trial omission (≤30 s of laser stimulation/trial).

### Behavioral control tests

#### Progressive ratio (PR) test

Following the final RDT session, mice that underwent fiber photometry and microendoscopic imaging were retrained to RM criterion and then underwent a single progressive ratio (PR) test session (modified from Heath et al. 2015). During PR, the large reward stimulus (alone) was presented in the same touchscreen response-window as the mouse had during RM discrimination. A touchscreen response at the large reward stimulus caused it to disappear for 1 second and then reappear. The first touch produced the large reward, reward tone, and illumination of the reward port. The number of touchscreen presses required for reward delivery increased by 4 after each reward (i.e., 4 touches for reward one, 8 touches for reward two, 12 touches for reward three, etc.)

A subset of optogenetic ChrimsonR and control mice underwent a single PR test (Laser On vs. Laser Off pseudorandomly assigned), conducted after retraining, as above. For mice assigned to the Laser On condition, optogenetic stimulation (same ChrimsonR parameters as above, lasting 5 s) was triggered after each touch of the large reward stimulus.

#### Footshock sensitivity

Mice received a footshock sensitivity test following the completion of touchscreen testing. Footshock was delivered in the touchscreen chamber every 10 seconds, ascending in intensity from 0.00 to 0.50 mA in 0.02 mA increments (i.e., 25 shocks total). In the case of chemogenetic inhibition (**Extended Data Fig. 2g**), CNO was delivered ∼30 min prior to the shock sensitivity session. In the case of stGtACR2-mediated inhibition (**Extended Data Fig. 5j**), laser illumination (same parameters as above) was timed to each shock delivery, and lasted 2 s.

Video of the mouse was analyzed offline by an experimenter blind to experimental treatment (if applicable) to score 3 responses: noticing (orienting to the grid floor and backwards treading), vocalization, and rapid escape (quick burst of forward motion), as previously described (Piantadosi et al. 2018). For correlational analyses (**Extended Data Fig. 1f**), the intensities corresponding to the first instance of noticing, vocalization, and rapid escape were summed to produce a ‘Footshock sensitivity’ measure.

### Data collection and analysis

#### Measurement of locomotion and choice-abort behavior

Video of touchscreen sessions were used to extract the position of the mouse within the touchscreen apparatus and quantify abort behavior (see below). For experiments involving quantification of neural activity (fiber photometry, microendoscopic imaging), a FLIR Blackfly S camera (BFS-PGE-16S2C-CS, Teledyne FLIR, Elkridge, MD, USA) was suspended above the chamber to record test-sessions. SpinView software (SpinView, Las Vegas, NV, USA) controlled the camera settings, and Bonsai software (Lopes et al. 2015) was used to acquire frames (30 FPS) and write these data to file. For microendoscopic imaging, video recording was triggered by the start of the behavioral program (after microendoscopic data acquisition had begun, see details below). For fiber photometry, video recording was triggered by the start of photometry data acquisition (see details below). These triggering approaches ensured the behavioral video was aligned with each activity measurement approach. For chemogenetic and optogenetic experiments, GoPro cameras (HERO 5 or HERO 7; GoPro, Clearview Way, San Mateo, CA, USA) were mounted above the chamber to acquire videos (30 FPS) after manual start. Camera frames were aligned to the behavioral session by manually labeling the first video frame in which the behavioral session began (indicated by house-light illumination), and adjusting all timestamps recorded within the touchscreen by the time interval between the start of the video and the start of the behavioral program.

Mouse position was quantified from video using the machine-learning algorithm Social LEAP Estimates Animal Poses (SLEAP) (Pereira et al. 2022). Random frames selected from a subset of videos were first manually labeled with 6 body points (microendoscopic imaging/fiber photometry: snout, left ear, right ear, neck, body, and tail base) or 5 body points (chemogenetics and optogenetics: snout, left ear, right ear, body, tail base). These frames were used to train the model iteratively following manual correction of errors prior to further labeling and re-training. Inference was then run on all unlabeled frames for all videos. The resulting positional data used to calculate movement-related variables (e.g., velocity) were filtered using the MATLAB function ‘sgolayfilt’ (order = 3, framelen = 25).

Aborted screen-touches during choice were manually scored by an experimenter (blind to experimental group, if applicable) using BORIS software (Friard and Gamba 2016), as previously described (Halladay et al. 2019). Aborts were defined as a stretched posture towards the screen followed by rapid retraction or shift in head direction away from the screen (**Supplemental Video S2**). To align aborts to neural activity, the video frame immediately preceding the retraction/head-shift was marked as the time of the abort.

#### Behavior data analysis

Within each block of trials, the percentage of choices of each option (Large^Rew^ and Small^Rew^) on free-choice trial was calculated as: (# of choices / 22) * 100. Choice latency was calculated as the time from trial initiation (IR beam break within reward receptacle) to choice across all trial types (forced & free-choice), in seconds. Reward collection latency was calculated as the time from choice to reward collection (IR beam break within reward receptacle) across all trial types (forced & free-choice), in seconds. Reward consumption latency was calculated as the time from reward collection start to reward collection end (the interval between the IR beam break triggering collection start, and the IR beam break triggered by the mouse’s head exiting the reward receptacle). Aborted choices were quantified as described above. To calculate locomotor path lengths, X and Y coordinates corresponding to the mouse ‘Body’ (defined by SLEAP label, as described above) point were extracted. The point corresponding to the body was used as it was the most consistently well-tracked point among all points labeled. To calculate path lengths for each trial, the MATLAB function ‘pdist2’ was first used to extract the Euclidean distance between each sample, after which the sum of the distances was calculated as the path length. Data related to these behavioral variables were analyzed using twoway repeated-measures ANOVAs with Choice Type and Trial Block as within-subjects factors, with Tukey’s test for multiple-comparisons.

PR measures were extracted including the final PR ratio (the last PR ratio achieved before session end) and the total number of presses made during the session. For the analysis of PR data from ChrimsonR vs. control mice (**Extended Data Fig. 10c**), a two-way ANOVA with Treatment (ChrimsonR vs control) and Laser Status (Laser On vs. Laser Off) as between-subjects factors was conducted, with Tukey’s test for multiple-comparisons.

To analyze the effect of chemogenetic (e.g., **Fig. 1k**) and optogenetic (e.g., **Fig. 4j & 6k**) manipulations on decision-making behavior, data were analyzed with two-way between/within-subjects ANOVAs, with Treatment as the between-subjects factor, and Trial Block as the within-subjects factor. Multiple comparisons were Tukey’s tests to compare Treatment groups for hM4Di vs mCherry and stGtACR2 vs Control, or Dunnett’s tests for ChrimsonR and PdCO vs Control.

To calculate the effect of risk on reward collection and trial initiation latencies, collection and initiation (defined as the interval between a completed trial and the initiation of the subsequent trial) latencies were calculated for each forced and free-choice trial. Then, a bootstrapped sample based on the safe trial block data (trials 1 to 30; number of bootstraps = 1000) was created using the MATLAB function ‘bootstrp’ and the confidence interval (CI) was calculated across these samples (alpha = 0.001). Trial-wise significance was indicated if the mean latency (across mice) exceeded the bootstrapped CI for a given trial.

A paired-samples t-test was used to evaluate the difference between the number of trials on which punishment was followed by an aborted choice (Abort^Large^ or Abort^Small^) versus a choice (Fig, 3i)

To analyze choice preference (Large^Rew^ vs Small^Rew^) across behavioral training (Early vs Late RM), a two-way repeated measures ANOVA with Choice Type and Training Stage was used, with multiple-comparisons conducted using Tukey’s tests. To analyze zone time within the trapezoidal touchscreen arena (**Extended Data Fig. 1c & 3c**), zones corresponding to Large^Rew^ (**l**) and Small^Rew^ (S) response-windows, reward port (R), and the rest of the arena (other, O) were manually drawn within the arena using custom-written MATLAB scripts. X and Y coordinates corresponding to the mouse’s body point (defined by SLEAP label, as described above) were extracted and used to calculate, for every trial, the proportion of time spent in each zone. These data were analyzed using a two-way repeated measures ANOVA, with Zone and Trial Block as within-subjects factors, with multiple-comparisons conducted using Tukey’s tests.

Correlations between Large^Rew^ choice and the variables weight or footshock sensitivity (**Extended Data Fig. 1d-f**) or final PR ratio and total presses (**Extended Data Fig. 1h-i**) were conducted using Pearson’s correlations (MATLAB function ‘corr’).

### Calcium imaging data analysis

#### Microendoscopic imaging and preprocessing

Data were collected as previously described (Silverstein et al. 2024) using an Inscopix data acquisition system connected to a laptop computer running Inscopix Data Acquisition Software (IDAS). Videos of Ca^2+^ dynamics were acquired at 20 Hz (50 FPS) using either the nVista, nVue1.0, or nVue2.0 (Inscopix) miniaturized 1-photon microendoscopes. The start of the behavioral session sent a transistor-transistor logic (TTL) pulse to trigger the video camera to record behavior (as described above) and to timestamp the start of the ABET II Touch behavioral program within IDAS. Ca^2+^ videos underwent initial preprocessing using custom-written Python scripts by accessing the Inscopix ISX API. Videos were first downsampled (4x spatial bin, 2x temporal bin), then spatially filtered (low cut-off = 0.005, high cut-off = 0.5), followed by motion correction, as previously described (Thevenaz et al. 1998). Preprocessed data were saved as .tiff files and custom-written MATLAB scripts (MathWorks, Natick, MA, USA) were used to identify putative single neurons by implementing a constrained non-negative matrix factorization algorithm optimized for endoscopic data (CNMFe, Zhou et al. 2018). Florescence data corresponding to these CNMFe-identified neurons was used for all subsequent analysis.

#### Cross-session registration of neurons

To track neurons across sessions (e.g., **Extended Data Fig. 4a,b**), CNMFe-identified cell outlines (footprints) from pairs of behavioral sessions were aligned using CellReg (Sheintuch et al. 2017). For most session pairs, the spatial correlation model was used for registration, though some sessions required the centroid distance to be used. The minimum criterion for a cell pair to be classified as being the same was a probability > 0.6. For a small number of session pairs, neuron footprints in one or both sessions were too sparse to align using CellReg (mostly BLA-NAcSh imaging sessions, where fewer neurons on average are seen within each FOV). In this case, FOVs were visually inspected and manually aligned using custom written MATLAB scripts.

#### Event-specific sub-window definitions

To define whether neurons were active during a given event or task epoch, sub-windows of interest were defined as follows: Pre-choice = -4 to 0 seconds relative to choice, Post-choice^Rew^ = 0 to +2 seconds relative to choice, Post-choice^Pun^ = 0 to +2 seconds relative to choice, Consum. = +1 to +3 seconds relative to reward collection, Abort = 0 to +2 seconds relative to abort onset (see definition above).

#### Event-related ensemble identification

Neurons were assigned to a specific ensemble based on a previously described method (Jimenez et al. 2018, Piantadosi et al. 2024a). For each event-specific category (e.g., Pre-choice, Post-choice^Rew^ or Post-choice^Pun^, Consum., Abort), Z-scored data for each trial (aligned to the event of interest) were extracted. The mean activity within the event-specific sub-window (see above) was calculated across all trials, forming the empirical mean for that event, for that neuron. Then, a shuffled distribution was created to compare with the empirical mean: the Z-scored empirical data were shuffled in time (for each trial) and a trial-mean was calculated, with this process repeated 1000 times, to create a distribution of event-aligned, shuffled data. A neuron was determined to be significantly active in response to a given event if its mean empirical response within the event sub-window exceeded the mean of the shuffled distribution by 1.5 SD. Due to the difficulty in determining the meaning of a negative deflection in ΔF/F due to the decay properties of each GECI used here, only neurons that displayed increases in fluorescence activity were included in a given category. Neurons were described as being exclusively responsive during Pre-choice, Post-choice^Rew^, or Consum. periods (e.g., **Fig. 2h**) by determining whether the activity in one period (e.g., Pre-choice) exceeded the threshold stated above, but did not for the other two categories (e.g., Post-choice^Rew^ or Consum. periods).

The above process was conducted separately for the initial trial block and for the combination of the second and third trial block. This allowed assessment of changes across each task (Late RM and RDT) driven by changes in contingency (e.g., the introduction of punishment risk). Because the number of trials for a given comparison was unbalanced by the combining of the second and third block, these trials were randomly sub-sampled to match the number of trials in the first block.

Once identities were determined, we calculated the sum and percentage of neurons within each category for the pseudopopulation of all recorded neurons (e.g., **Fig. 2g**). Statistical comparison of the proportions of different populations (e.g., **Fig. 3g**) was conducted using a Chi-square test of proportions (via custom written MATLAB scripts). To compare the relative sizes of identified ensembles for projection-specific (BLA^→^NAcSh) and pan-neuronal imaging (pan-BLA) groups (**Fig. 6g,h**), two-proportion Z-tests were used (conducted using custom-written MATLAB code). The proportion of neurons that were Conserved, Lost, or New during early versus late trial blocks for Late RM and RDT (**Extended Data. Fig. 5**) were compared using Chi-square test of proportions (via custom written MATLAB scripts).

To describe ensemble dynamics during safe reward-seeking for Large^Rew^ and Small^Rew^ trials (**Extended Data Fig. 3e**), ensembles were identified (as above) for Small^Rew^ trials only. Because statistical assignment of neurons to particular ensembles is dependent on the number of trials, the same process was repeated using data from Large^Rew^ trials, randomly downsampled to the number of trials present for the Small^Rew^.

#### K-means clustering

Unbiased K-means clustering of neurons based on activity (e.g., **Fig. 2e**) was conducted in a manner similar to Howe et al. 2024. Fluorescence data from Pre-choice, Post-choice^Rew^, and Consum. neurons were aligned to choice, Z-scored, and then the mean within each event-specific sub-window was calculated. This resulted in each neuron being represented by 3 values corresponding to its mean response within each epoch. The resulting n-neuron x 3 array was used for K-means clustering using the ‘kmeans’ function in MATLAB, with the distance metric being ‘correlation’, and number of clusters = 3.

#### Decoding

Decoding was performed within each mouse, largely as previously described (Silverstein et al. 2024). For decoding neuron identity during safe reward-seeking (**Fig. 2f**), fluorescence data aligned to choice (for Pre-choice and Post-choice^Rew^ neurons) or consumption (for Consum. neurons) was extracted for each neuron. Then, an identical array was created by extracting data in the same way as above, but from neural data that had been circularly rotated to temporally desynchronize the activity from the event timing. Empirical data and its shuffled counterpart from neurons that were identified as Pre-choice, Post-choice^Rew^, or Consum. were then reduced in time by calculating the mean activity for each neuron within the relevant event-specific sub-window (see above for definition). Thus, for every neuron, the array to be decoded was made up of one array that reflected the empirical data (size = # of events x 1), and another identically-sized array reflecting the shuffle. Then, for each mouse, these arrays were vertically concatenated and Z-scored, assigned values of 0 (empirical data) or 1 (shuffle), and partitioned into test and training datasets using the MATLAB function ‘cvpartition’ with 5-fold cross-validation (KFold = 5). Decoding was conducted using a support vector machine (SVM) classifier via the MATLAB function ‘fitcsvm’. Decoder performance was evaluated within each mouse from the mean F1 score across the 5 cross-validation folds (empirical data compared to the shuffle). These mean F1 scores were compared to mean F1 scores generated by directly decoding two shuffled arrays (generation of shuffled arrays and decoding conducted as above) using paired t-tests (conducted using the MATLAB function ‘ttest’).

#### Correlations

Pairwise correlations between neurons in each ensemble (e.g., **Fig. 2i**) were conducted by first vertically concatenating the mean activity (across all trials) for each neuron in the pseudopopulation. Then, each row was correlated with every other row (pairwise correlations of mean trial-wise activity for each pair of neurons). The resulting correlations were then split into within- and across-categories for every pair of neurons to calculate the proportion of positive, negative, and neutral correlations across each possible pairwise group comparison.

For correlations between trial-wise behavioral variables and neural activity (e.g., **Fig. 2k**), mean Z-scored fluorescence activity for every neuron was calculated within the epoch-specific sub-window of interest (e.g., Pre-choice, -4 to 0 s from choice) on a trial x trial basis, resulting in a 1 x # of trials array. For each neuron, this array was correlated (using the MATLAB function ‘corr’) with the behavioral variable of interest (e.g., trial choice latencies, which is a 1 x # of trials array). These resulting correlations could then be broken down across ensembles, such that the correlations of Pre-choice neurons with trial choice latency (for example) could be compared to correlations conducted in the same fashion but from non-responsive neurons. Distributions of correlations for the ensemble of interest versus non-responsive neurons were compared using Kolmogorov-Smirnov tests using the MATLAB function ‘ktest2’. Mean correlation coefficients were compared using unpaired t-tests using the MATLAB function ‘ttest2’.

Correlations between the proportion of ‘New’ neurons of a given ensemble identity and risk preference (% large choice) (e.g., Fig 4f) were conducted using Pearson correlations (MATLAB function ‘corr’).

#### Dimensionality reduction using principal component analysis

To conduct dimensionality reduction as a method to investigate population-level representations (e.g., **Fig. 3h**), mean fluorescence data was collected across trials in a window aligned to each event of interest. These data were then horizontally concatenated (across events) and Z-scored before conducting dimensionality reduction using principal component analysis (PCA) via the ‘pca’ MATLAB function. Neural data were then projected through time for the two PCs explaining the most variance by multiplying the PCA coefficients/loadings by the fluorescence data separately for each event analyzed.

To statistically compare PCA trajectories across two events, the Euclidean distance (in arbitrary units) for PC1 was calculated at each timepoint. In addition, a null distribution was created by conducting the same PCA procedure on circularly rotated data 100 times. The Euclidean distances generated from the empirical data were directly compared to the bootstrapped null distribution, such that time periods where the empirical data diverged from the null distribution (alpha = 0.001) were deemed significant.

#### Permutation tests for timeseries statistical comparison

To assess whether time-varying neural signals differed significantly (e.g., **Fig. 3f**), permutation tests and confidence-interval based approaches were used (Jean-Richard-dit Bressel et al. 2020). For permutation tests, Z-scored mean data (across trials) for each sample were vertically concatenated and randomly resampled (neurons shuffled to new row indices) before taking the difference between the means of the randomly resampled data. Random resampling and differencing was repeated 1000 times. For every time bin, we calculated the proportion of random resamples that had a larger absolute difference than the observed data, with consecutive bins (minimum consecutive temporal bins = 10) where the proportion was lower than the alpha level (0.001) deemed significantly different.

To assess whether the change in ΔF/F between safe and risky blocks was significant (**Fig. 4b**), we tested whether confidence intervals surrounding the difference score overlapped with 0. Z-scored mean data (across trials) for neurons from each ensemble identified during the safe block (e.g., Pre-choice) were extracted for both the safe block and the risky block. The difference between these matrices was calculated and used as the input for the bootstrapped CI (bCI) testing. The empirical difference array was resampled and the mean calculated 1000 times to create a bootstrap distribution from which an upper and lower CI could be generated (alpha level = 0.0001). Samples where the upper CI was greater than 0, or the lower CI was less than 0 for a minimum of 10 consecutive temporal bins were deemed statistically significant.

#### Population vector correlations

Population vector (PV) correlations representing particular epochs or events were used to examine neural population-level representations (Courtin et al. 2022, Rozeske et al. 2023). PVs were assembled (within each mouse) by first taking the mean activity within the epoch-specific sub-window (e.g., Pre-choice: -4 to 0 around choice) on each relevant trial. This resulted in every mouse having a # of neurons x # of trials matrix, from which the mean was calculated across trials. The resulting PV was of the size # of neurons x 1 (mean activity). This vector was correlated with session-long neural activity (100 ms bins), such that each mouse had a PV correlation reflecting the ensemble or event relatedness across time.

For analyses where PV correlations were compared across safe and risky trial blocks (**Fig. 4d**), PV correlations were calculated as above, separately for ensembles identified during the safe trial block or the risky trial blocks. Then, for each trial and for each ensemble, the mean PV correlations within the relevant epoch-specific sub-window was extracted. The mean of the sub-window extracted PV correlations were calculated for trials within the safe block and the risky trial blocks, with safe-identified and risky-identified data compared using paired-samples t-tests within safe and risky blocks using the MATLAB function ‘ttest’.

### Fiber photometry analysis

#### Fiber photometry

Data were collected as previously described (Gunduz-Cinar et al. 2023, Sengupta and Holmes 2019) on a computer interfacing with a RZ5P processor (Tucker-Davis Technologies, Alachua, FL, USA). Illumination from 465 nm and 405 nm light-emitting diodes (LEDs; Thorlabs) were modulated at non-resonant carrier frequencies and passed through a fluorescence minicube (Doric Lenses). Modulated light passed through a fiber optic rotary joint (Doric Lenses, catalog#: FRJ_1x1_PT_400-0.57_2m_FCM_0.12m_FCM) that was attached to a fiber-optic patch cable (Doric Lenses, catalog#: MF_400/430/LWMJ-0.57_0.5m_FCMMF1.25(**f**). The patch cable was attached to the fiber-optic implant via a mating sleeve (Doric Lenses, catalog#: SLEEVE_ZR_1.25). Emitted fluorescence was sampled at 6.1 kHz and detected by a photoreceiver (Newport Corporation, Irvine, CA, USA). Demodulated 405 nm (ultra-violet control channel) and 465 nm (GCaMP6m emission) wavelength light were saved for offline analysis. Data from each wavelength were lowpass-filtered and downsampled.

Data corresponding to the 465 and 405 channels were extracted around a 10-second peri-choice period for each trial and the 405 nm signal was scaled using a linear least-squares regression to fit the 465 nm signal. Event-related change in fluorescence (ΔF) was calculated as 470 nm – fitted 405 nm signal. ΔF data for each event was Z-normalized by subtracting the mean of the ΔF signal within the event window and dividing this by the standard deviation of the event window ΔF signal. Periods in which Z-normalized peri-event activity differed from null were statistically determined as 95% confidence intervals (bCI) calculated from a 1000-fold bootstrap estimate, as previously described (Jean-Richard-dit Bressel et al. 2020). For quantifying the degree of pre-choice activity, area under the curves (AUCs) during the pre-choice period (-4 to 0 s before choice) were calculated for safe versus risky blocks (RDT) or for the initial block and the final two blocks (Late RM) for each mouse. These data were statistically compared using paired t-tests, conducted using the MATLAB function ‘ttest’.

### Ex vivo procedures

#### General histology procedures

Mice were injected with a terminal dose of FatalPlus (pentobarbital) and transcardially perfused with ice cold phosphate-buffered saline (0.2M PBS), followed by a 4% paraformaldehyde solution (PFA). For mice with head-mounted optical interfaces, the entire head was submerged in 4% PFA overnight at 4°C, before removal (to allow for accurate visualization of fiber or lens depth) and a second overnight 4% PFA fixation. Brains were then transferred to a 0.1M phosphate buffer solution and stored at 4°C until sectioned. Unless otherwise specified, 50-µm serial coronal sections were prepared using a vibratome (Model VT100S, Leica Biosystems, Buffalo Grove, IL, USA), with every third section mounted directly on to ColorFrost Plus microscope slides (Fisherbrand; 12-550-20). Hoechst (20 mM; Thermo Scientific; 62249) was diluted in 1X Dulbecco’s PBS (Gibco) to a final concentration of 0.0163 mM and slides containing tissue were submerged for 3 minutes before being coverslipped with a Mowiol mounting solution. Brain section-containing slides were scanned using an VS120 Virtual Slide Scanner (Olympus, Center Valley, PA, USA) at 10x magnification (PLAN APO, 0.4 numerical aperture, Leica). Atlas plates for plotting viral expression and optical implants (e.g., **Extended Data Fig. 1d**) were adapted from Paxinos and Franklin (2019).

#### Brain Street View (BSV) analysis

Projection densities were visualized with Brain Street View (BSV; https://github.com/Julie-Fabre/brain_street_view), using data from (Oh et al. 2014). Experiments where an EGFP-containing anterograde tracing virus was injected into BLA in C57BL/6J mice were included (n=3, experiment IDs: 277710753, 113144533, 120282354). Fluorescence values were normalized to the fluorescence intensity of the injection site (BLA) for each brain.

#### BLA output tracing quantification

For quantification of overlap of BLA output neurons projecting to NAcSh versus PL, tissue sections were first manually aligned to the Allen Mouse Brain Atlas. Neurons within the BLA were identified manually within OlyVIA software (V3.1.1, Olympus). Counts for GFP, tdTomato, and co-expressing neurons were recorded for each section containing the anterior-posterior extent of the BLA (sections separated by ∼200 µm). To examine the anterior-posterior gradient, counts were summed across the following ranges (distance in mm from bregma), Anterior: -1.06 to -1.58; Mid: -1.70 to -2.18; Posterior: -2.3 to -2.8. Data were analyzed using a repeated measures ANOVA with Pathway (BLA^→^NAcSh, BLA^→^PL, and Co-expressed) and Location (Anterior, Mid, Posterior) as within-subjects factors, with multiple comparison tests conducted using Tukey’s tests. Analyses were conducted using custom-written MATLAB code.

#### cTRIO rabies tracing quantification

For rabies quantification, tissue sections were manually aligned to the Allen Mouse Brain Atlas and brain regions containing fluorescently labeled neurons were outlined within OlyVIA software (V3.1.1, Olympus). GFP+/Rabies, mCherry+ and co-expressing neurons were manually counted for each brain region throughout the brain and GFP+/Rabies counts for each region were normalized to the total number (brain-wide) of GFP+/Rabies neurons. Only regions wherein at least one brain had >1% of rabies-positive cells are included. Counts for regions where starters & rabies were injected (IL, BLA, and vHPC) and proximal regions where >10 starter cells were found in at least one brain are not included for each pathway. Brain region naming and abbreviations follow the Allen Mouse Brain Common Coordinate Framework (CCFv3), with minor alterations. Because regions were omitted for some brains and not others (see description above), data were analyzed using unpaired t-tests with Welch correction (conducted within the statistical software GraphPad).

#### TRAPseq tissue processing

Mice were rapidly decapitated, brains were removed and rinsed in ice-cold Buffer B (1X HBSS, 4 mM NaHCO3, 2.5 mM HEPES, 35 mM glucose, 100 µg/ml cycloheximide). A ∼1.5 mm coronal section roughly spanning the anterior-posterior length of the BLA was cut using 2 stainless-steel blades, and a 1 mm tissue punch of each (bilateral) BLA taken and briefly stored in an aliquot containing ice-cold Buffer B (as above). Samples were processed in triplicate, such that bilateral punches from 3 mice were acquired, stored in ice-cold Buffer B, then transferred to an ice-cold Teflon/glass homogenizer (details) containing ∼1 ml of Buffer C (10 mM HEPES, 150 mM KCl, 5 mM MgCl2, 0.5 mM DL-Dithiothreitol (DTT), 80 U/ml RNasin Plus Ribonuclease Inhibitor (Promega), 40 U/ml SUPERase-In RNase Inhibitor (Invitrogen, Waltham, MA, USA), 100 µg/ml cycloheximide, protease inhibitor cocktail (Roche cOmplete, Sigma-Aldrich, St. Louis, MO, USA). Tissue in Buffer C were hand-homogenized on ice (10 strokes), followed by high-speed homogenization at ∼750 rpm (Glas-Col). The sample was then transferred to a new ice-cold aliquot and centrifuged at 2,000 xg for 10 minutes at 4°C. The supernatant was collected, and 1/9th volume of 10% IGEPAL CA-630 (NP-40 replacement; SupeIco, Sigma-Aldrich) was added (mixed gently by inversion, and spun on a minifuge), followed by 1/9th volume of DHPC (07:0 PC; Avanti Polar Lipids, Alabaster, AL, USA), before being incubated on ice for 5 minutes. The solution was mixed by inversion and centrifuged at 17,000 xg for 15 minutes at 4°C.

The supernatant was transferred to a new tube and used for immunoprecipitation (50 µl of the lysate was reserved and combined with 50 µl of lysis buffer (Buffer RLT from the Qiagen RNeasy Micro Kit, with 10 µl/ml beta-mercaptoethanol) and stored at -80°C for later analysis as input RNA). Magnetic beads (300 µl; Dynabeads MyOne Streptavidin T1; Invitrogen) were washed twice on a magnetic rack with 1x PBS, and then incubated in a 1x PBS solution with 120 µl of biotinylated Pierce Recombinant Protein L (Thermo Scientific) for 35 minutes on a tube rotator at room temperature. Protein L loaded beads were blocked with five washes of 1x PBS containing 3% IgG-free and protease-free Bovine Serum Albumin (BSA; Jackson ImmunoResearch). After the final block, beads were suspended in Buffer A (10 mM HEPES [pH 7.4], 150 mM KCl, 5 mM MgCl2, 1% IGEPAL CA-630), loaded with anti-GFP antibodies (Memorial Sloan Kettering Cancer Center, Bioreactor Supernatant quality) 19C8 (50 µg; HtzGFP-19C8) and 19F7 (50 µg; HtzGFP-19F7), and spun on a tube rotator at room temperature for 1 hour.

Beads containing anti-GFP antibodies were washed twice in Buffer A (plus 0.5 mM DTT, 40 U/ml RNasin Plus, and 100 µg/ml cycloheximide), resuspended in Buffer A and combined with brain lysates before incubating on a tube rotator at 4°C for 40 minutes. The bead and lysate mixture was then washed three times on ice with Buffer D (20 mM HEPES, 350 mM KCl, 10 mM MgCl2, 1% IGEPAL CA-630, 0.5 mM DTT, 80 U/ml RNasin Plus, 100 µg/ml cycloheximide). The mixture was transferred to a new, ice-cold aliquot, washed once more with Buffer D, after which the buffer was removed, the tube warmed to room temperature, and the RNA was eluted by adding 100 µl lysis buffer (Buffer RLT from the Qiagen RNeasy Micro Kit, with 10 µl/ml beta-mercaptoethanol). This mixture was vortexed and incubated at for 10 minutes at room temperature, after which RNA (now in lysis buffer) was moved to a new tube and purified using the Qiagen RNeasy Micro Kit (Qiagen, Hilden, Germany). These samples were then sent for sequencing at the Functional Genomics Core at UCSF.

#### TRAPseq data analysis

Sequencing was conducted at the Functional Genomics Core at UCSF, largely as in Gergues et al. (2020). Briefly, sequencing library construction was conducted using SMART-seq.v4 RNA kits (Takara Bio, Mountain View, CA, USA) and Nextera XT DNA Library preparation kits with multiplexing primers (Illumina, San Diego, CA, USA). Quality control (fragment quality, size, and concentration) was conducted using a 5200 Fragment Analyzer (Agilent, Santa Clara, CA, USA). Libraries were multiplexed at seven per flow-cell lane and sequenced on a HiSeq 4000 SE50 (Illumina). Transcript reads were aligned to the mouse genome (GRm38) using STAR 2.7.2b.

Raw count data were normalized using variance-stabilizing transformation (VST), and outlier samples were identified iteratively based on Mahalanobis distances in principal component space (PC1 and PC2). A chi-squared threshold (p < 0.01) was used to flag extreme observations, which were removed in successive rounds until no further outliers were detected. A total of four samples were excluded: vmPFC.4, vmPFC.3, PL.2 and PL.3 (**Extended Data Fig. 7b**). Lowly expressed genes were filtered using filterByExpr from the edgeR package, retaining only those expressed in at least 70% of samples (min.prop = 0.7). Differential expression analysis was performed using DESeq2’s likelihood ratio test (LRT) to assess group-wise differences across all projection-defined populations. In parallel, one-vs-rest DESeq2 models were applied to identify region-specific marker genes by contrasting each projection group against all others. For both approaches, differentially expressed genes (DEGs) were defined by a false discovery rate (FDR) < 0.1 (Benjamini-Hochberg correction) and an absolute log2 fold change exceeding 0.263, corresponding to a ≥20% change in expression. Fold changes were estimated using the lfcShrink() function with the “ashr” method to improve interpretability and reduce variance in lowly expressed genes. NAc-specific DEGs were further analyzed for functional enrichment using Gene Ontology (GO) and KEGG pathway analysis via the enrichR package. To identify minimal gene sets capable of distinguishing projection-defined subtypes, we applied sparse Partial Least Squares Discriminant Analysis (sPLS-DA) using the mixOmics R package. A one-vs-rest classification strategy was used for each projection group, with model parameters optimized through repeated cross-validation. Overlap between DESeq2-derived DEGs and sPLS-DA-selected features was quantified to evaluate concordance between univariate and multivariate feature sets.

## Abbreviations

RDT: risky deicison-making
BLA: basolateral amygdala
NAcSh: nucleus accumbens shell

## AUTHOR CONTRIBUTIONS

P.T.P. and A.H. conceived of and designed each experiment. P.T.P., K.M.C., H.C., S.J.P., M.H., J.A.S., J.P.G., N.A.S., N.R.S., R.S.V., S.S., and M.M.D. performed experiments. P.T.P., R.S., and D.S. analyzed data. P.T.P. and A.H. wrote and edited the manuscript. A.H. supervised the project.

## ACKNOWLEDGMENTS

The authors would like to thank Ruairi O’Sullivan for his advice regarding data analysis, Olena Bukalo and Ozge Gunduz-Cinar for their advice and technical assistance, Brian Copits for generously providing virus, and Lenka Maliksova and other members of the Functional Genomics Core at University of California San Francisco for their assistance with sequencing.

## DATA AVAILABILITY

Data available from the corresponding authors on reasonable request.

## SUPPORTING INFORMATION

Additional supporting information may be found in the online version of the article.

## SUPPLEMENTAL FIGURES

**Extended Data Figure 1:**
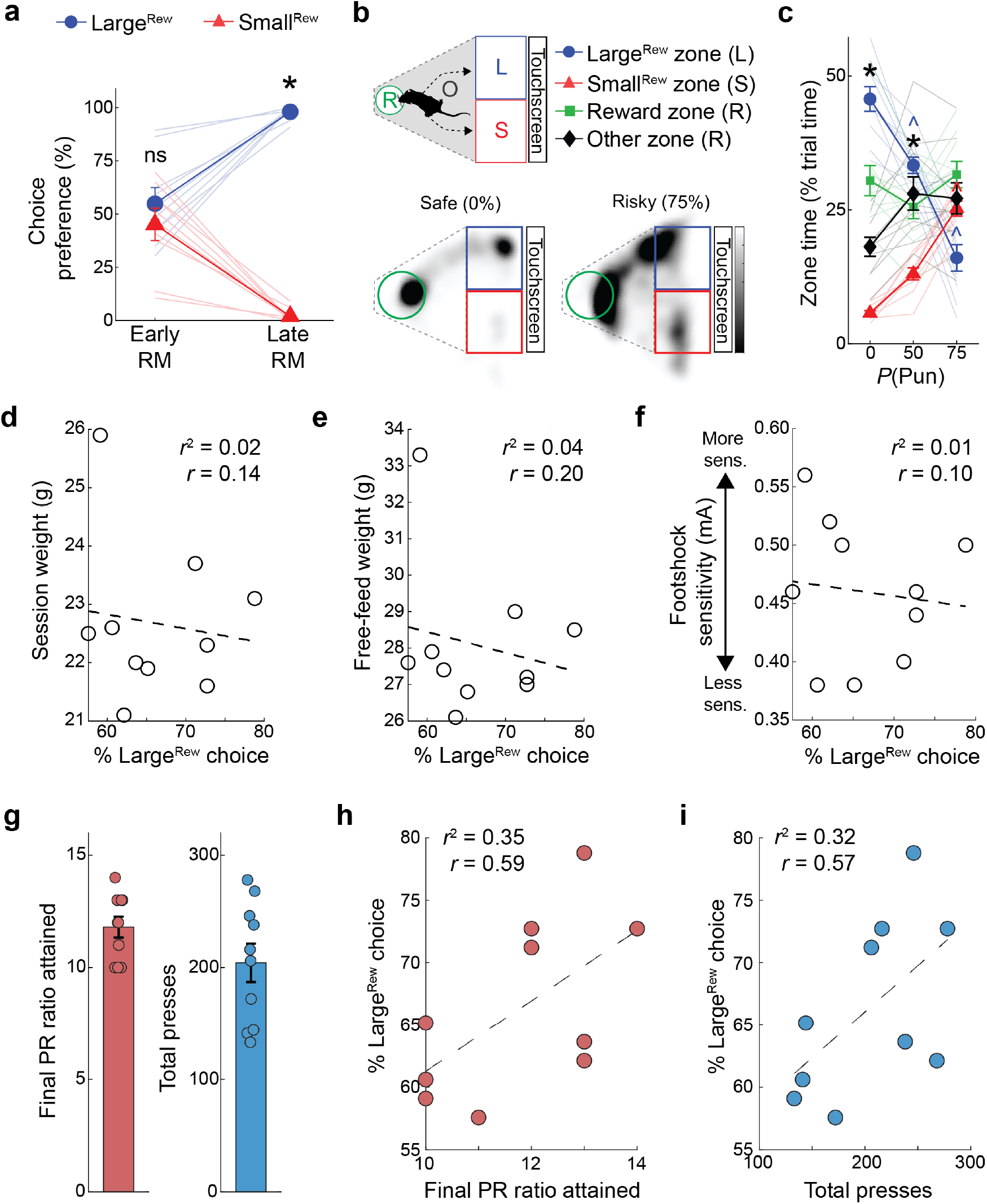
Behavioral measures during various behavioral tests. (**a**) Higher preference for Large^Rew^ versus Small^Rew^ on Late (p<0.001; ANOVA interaction, F(1,9)=45.68, p<0.001) not Early RM. (**b**) Upper: Cartoon depicting zones within touchscreen chamber corresponding to Large^Rew^ (L) and Small^Rew^ (S) response-windows, reward port (R), and the rest of the arena (other, O). Lower: Representative heatmap of time-spent during safe and 75% risky block. (**c**) Punishment caused mice to alter the amount of time spent within distinct touchscreen chamber zones (ANOVA interaction, F(6,54)=39.37, p<0.0001). Time spent in the Large^Rew^ zone decreased during risky blocks versus the safe block (within Trial Block effect for Large^Rew^: p<0.01 for both risky blocks vs safe block), while time in the Small^Rew^ zone increased (within Trial Block effect for Small^Rew^: p<0.01 for 75% risky vs safe block). Time in the reward and other zones did not significantly change across blocks. (**d-f**) Neither weight nor footshock sensitivity significantly predicted Large^Rew^ choice during the RDT. (**g**) Response measures in a touchscreen-based progressive ratio (PR) test. (**h,i**) Large^Rew^ preference was not correlated with final PR ratio (**h**) or total PR presses (**i**). n=10 mice for all comparisons. Multiple comparisons (Tukey’s post-hoc tests) were conducted following significant Training Stage (Early RM & Late RM) x Choice Type (Large^Rew &^ Small^Rew^) or Trial Block x Zone (Large^Rew^, Small^Rew^, Reward, Other zones) ANOVA interactions. For full reporting of statistics, see **Supplementary Table 1**). *p<0.05 other choice type (**a, c**). ^p<0.05 versus safe block, color corresponding to choice type (**c**). Data shown as mean ± SEM.

**Extended Data Figure 2:**
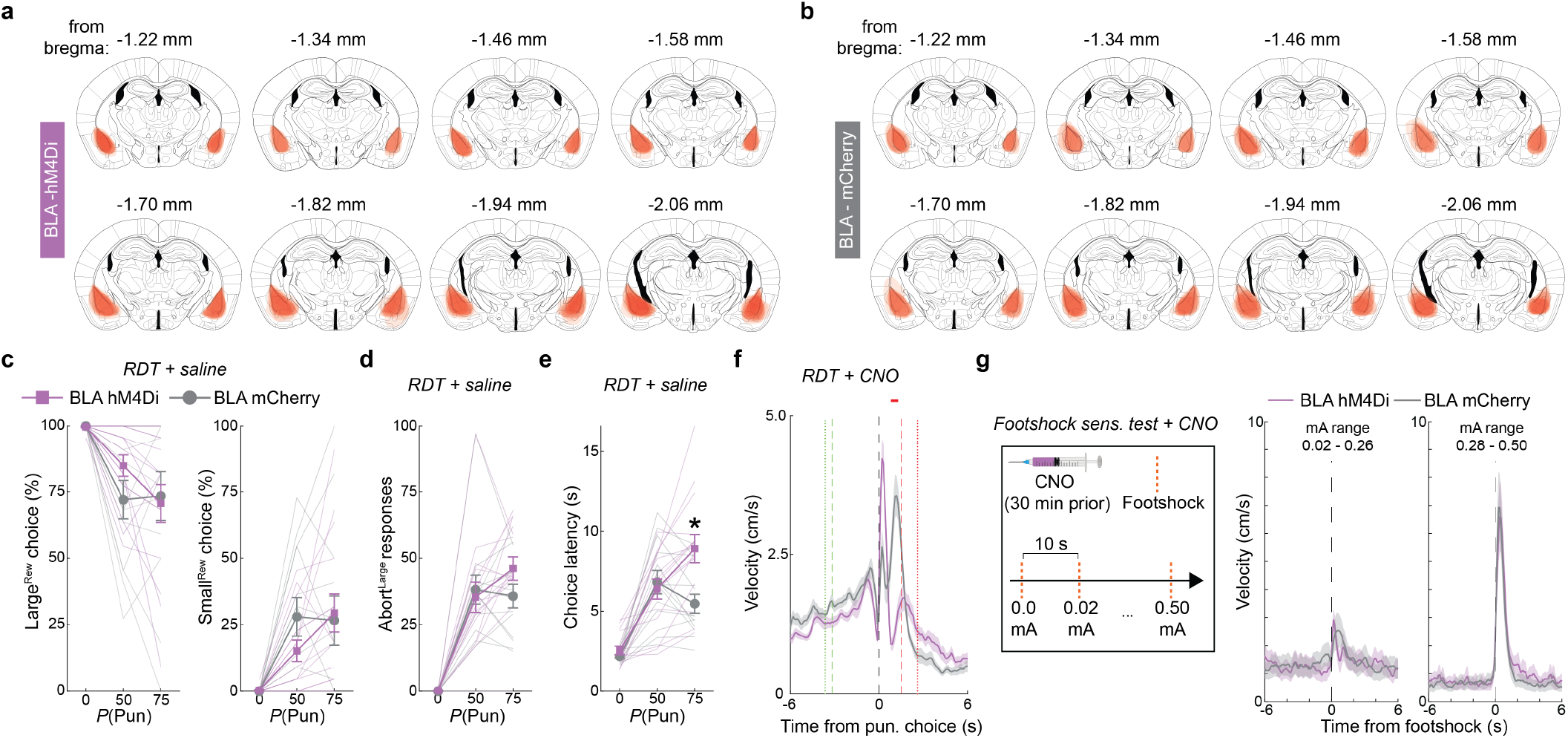
Chemogenetic BLA inhibition during RDT. (**a, b**) hM4Di (**a**) and mCherry control fluorophore (**b**) expression across the anterior-posterior extent of BLA. (**c-d**) Similar Large^Rew^ (left) and Small^Rew^ (right) choice (**c**) and Abort^Large^ responses (**d**) in hM4Di (n=15) and mCherry control (n=13) groups after saline injection. (**e**) Longer choice latency in hM4Di versus controls on the 75% risky block (hM4Di vs mCherry: p<0.01 for 75% risky block; ANOVA interaction: F(2,52)=9.27, p<0.005). (**f**) Slower movement velocity in response to footshock-punishment during RDT in the hM4Di versus mCherry controls (red horizontal line indicates group difference at corresponding samples, p<0.001, permutation test). Trial initiation times (green vertical lines) and reward collection times (red vertical lines) for hM4Di (dashed) and control (dotted) groups. (**g**) Schematic of footshock sensitivity assay following CNO administration (left). Similar footshock-induced movement velocity in hM4Di mice versus controls in an independent footshock sensitivity test at low (center) and high (right) shock intensity ranges. Multiple comparisons (Tukey’s post-hoc test) were conducted following significant Trial Block (0, 50, 75%) x Treatment (hM4Di & mCherry) ANOVA interaction. *p<0.05 versus mCherry. Data shown as mean ± SEM. Syringe (**g**, 10.5281/zenodo.4152947) graphic adapted from scidraw.io.

**Extended Data Figure 3:**
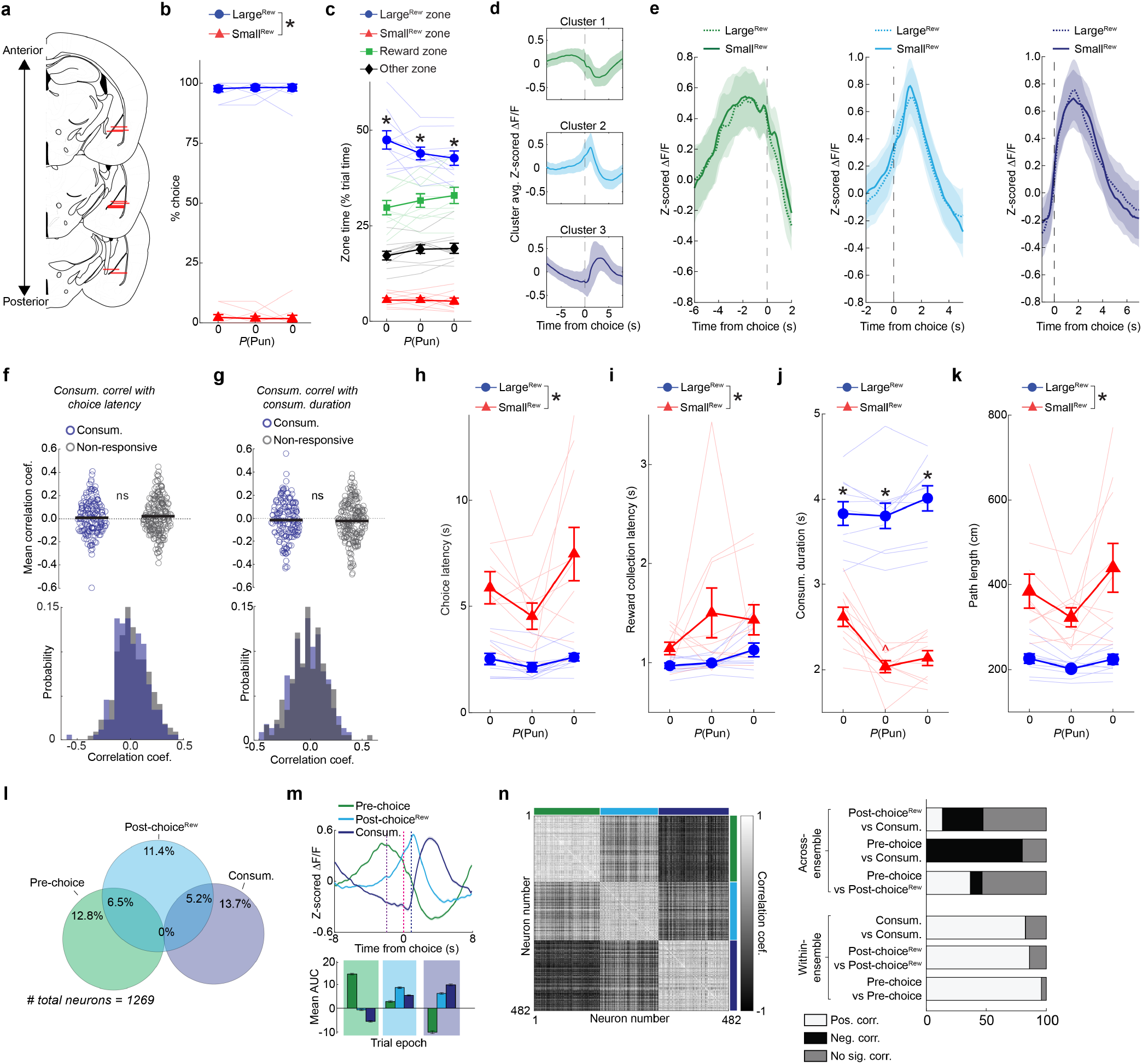
Behavioral and neuronal measures on reward magnitude (RM) test. (**a**) Ventral extent of GRIN lens placement (red lines) in BLA. (**b**) Near exclusive Large^Rew^ preference across trial-blocks on final reward magnitude session before RDT (Late RM) (F(1,9)=2373.76, p<0.001, repeated measures ANOVA, main effect of Choice Type). (**c**) More time spent in Large^Rew^ zone regardless of trial-block during Late RM (Large^Rew^ vs Small^Rew^: p<0.001 for all blocks; ANOVA interaction, F(6,54)=3.51, p<0.05) (see **Extended Data Fig. 1b** for zone definitions). (**d**) Mean neuronal activity for three K-means clusters analogous to Pre-choice (upper), Post-choice^Rew^ (center), and Consum. (lower) ensembles. (**e**) Indistinguishable choice-aligned activity of Pre-choice (left), Post-choice^Rew^ (center), and Consum. (right) ensembles identified from only Large^Rew^ or Small^Rew^ trials (trial-number matched across choice-types). (**f**) The activity of Consum. neurons was not related to choice latency. (**g**) Duration of reward consumption was not related to Consum. neuron activity. (**h,i**) Longer choice (**h**) and reward collection latencies (**i**) for Small^Rew^ versus Large^Rew^ (both F>15 and both p<0.001, repeated measures ANOVAs, main effect of Choice Type). (**j**) Longer reward consumption duration for Large^Rew^ versus Small^Rew^ on all Late RM blocks (Large^Rew^ vs Small^Rew^: p<0.001 for all blocks; ANOVA interaction: F(2,18)=14.99, p<0.001). (**k**) Longer path lengths for Small^Rew^ versus Large^Rew^ (F(1,9)=26.97, p<0.001, repeated measures ANOVA, main effect of Choice Type). (**l**) Degree of overlap between Pre-choice, Post-choice^Rew^ and Consum. ensembles. (**m**) Ensemble activity aligned to choice, with median trial initiation time (purple dashed line), time of choice (pink dashed line), and the median time of reward collection (dark blue dashed line) indicated (top); epoch ensemble AUCs (bottom). (**n**) Pair-wise correlation matrix organized by ensemble identity (indicated by colored bars) (left). Summary of across and within-ensemble correlations (right). Multiple comparisons (Tukey’s post-hoc test) were conducted following significant Trial Block (0, 50, 75%) x Choice Type (Large^Rew &^ Small^Rew^) or Trial Block x Zone (Large^Rew^, Small^Rew^, Reward, Other zones) ANOVA interaction. For full reporting of statistics, see **Supplementary Table 1**. Data mean ± SEM. *p<0.05 versus other choice type.

**Extended Data Figure 4:**
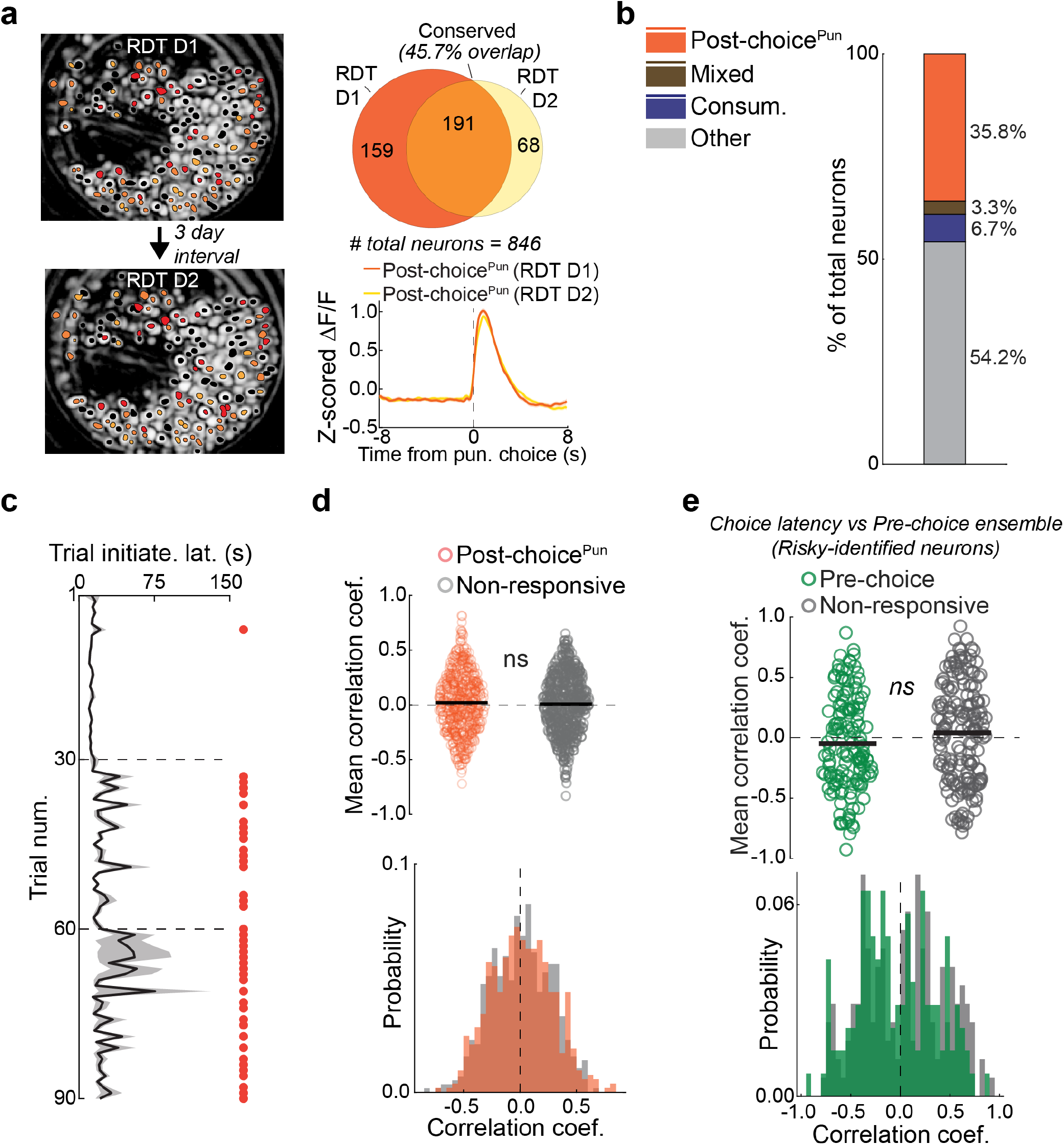
Cross-session activity of BLA ensembles and correlations with latency. (**a**) Imaging fields of view from two RDT sessions, conducted 3 days apart, with CNMF-E identified footprints color-coded by responsivity to shock on RDT D1 (red), RDT D2 (yellow), both (orange), or neither (black) (left). Many Post-choice^Pun^ neurons were recruited stably across days (upper right), with comparable activity profiles (bottom right). (**b**) Proportion of cross-session registered neurons encoding Post-choice^Pun^ during RDT, Mixed (responsive during consumption events on a progressive ratio test and during footshock delivery on the RDT), and Consum. (only responsive during consumption events on a progressive ratio test). (**c**) Longer trial initiation latencies during risky trials (trial blocks separated by horizontal dashed lines, red circles indicate trials where p<0.001, bootstrapped confidence interval computed from mean initiation latencies in the safe block). (**d**) Post-choice^Pun^ neuron activity during risky blocks was not related to trial initiation latency. (**e**) Pre-choice neuron activity during risky trial blocks was not predictive of choice latency. For full reporting of statistics, see **Supplementary Table 1**. Data mean ± SEM.

**Extended Data Figure 5:**
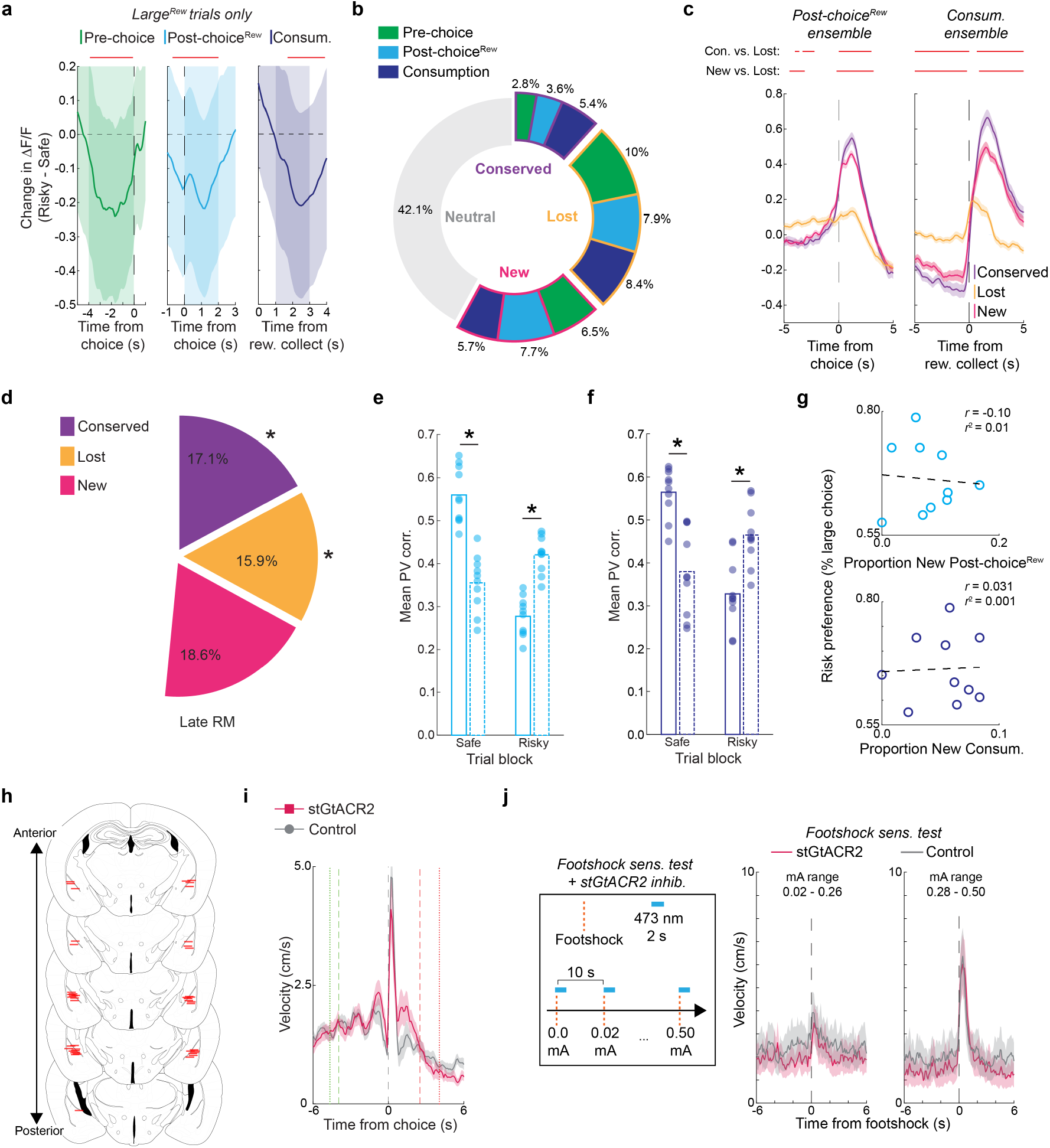
Additional risk-related BLA ensemble analyses and stGtACR2-based inhibition effects. (**a**) When examining data from Large^Rew^ trials only, decreased activity of Safe-identified ensembles during risky trial-blocks was still present (analysis as in **Fig. 4b**, red line indicates difference from 0, p < 0.001, bootstrapped confidence interval). (**b**) Proportion of neurons exhibiting Conserved, Lost, or New ensemble identity across Safe and Risky blocks for each ensemble category. (**c**) Mean activity traces for Post-choice^Rew^ (left) and Consum (right) ensemble neurons that were Conserved, Lost, or New (red horizonal line signifies samples different between groups indicated at left, p<0.001, permutation tests). (**d**) Significantly more Conserved (*χ*^2^(1)=10.36, p<0.001, Chi-square test) and fewer Lost (*χ*^2^(1)=36.07, p<0.001, Chi-square test) neurons during Late RM, as compared to RDT D1 (see **Fig. 4c**). (**e**) Mean PV correlations for Safe-identified and Risky-identified Post-choice^Rew^ ensemble neurons. Correlations for Safe-identified neurons were maximal on the safe block and declined on risky blocks (p<0.001, paired t-test), with the opposite being true of Risky-identified neurons (p<0.001, paired t-test). (**f**) As in **e**, for Consum. ensemble neurons (all p<0.001, paired t-tests). (**g**) Non-significant correlations between the proportion of New Post-choice^Rew^ neurons and risk preference (upper) and the proportion of New Consum. neurons and risk preference (lower). (**h**) Coronal mouse brain atlas sections depicting ventral extent of bilateral optic fibers (red lines) in BLA. (**i**) Similar movement velocity in stGtACR2 and control groups in response to footshock punishment during RDT. (**j**) Schematic depicting 473 laser delivery during footshock (left). stGtACR2 and control mice did not differ in response to footshock delivery at low intensity (center) and high intensity (right) ranges. For full reporting of statistics, see **Supplementary Table 1**. Data mean ± SEM. *p<0.05

**Extended Data Figure 6:**
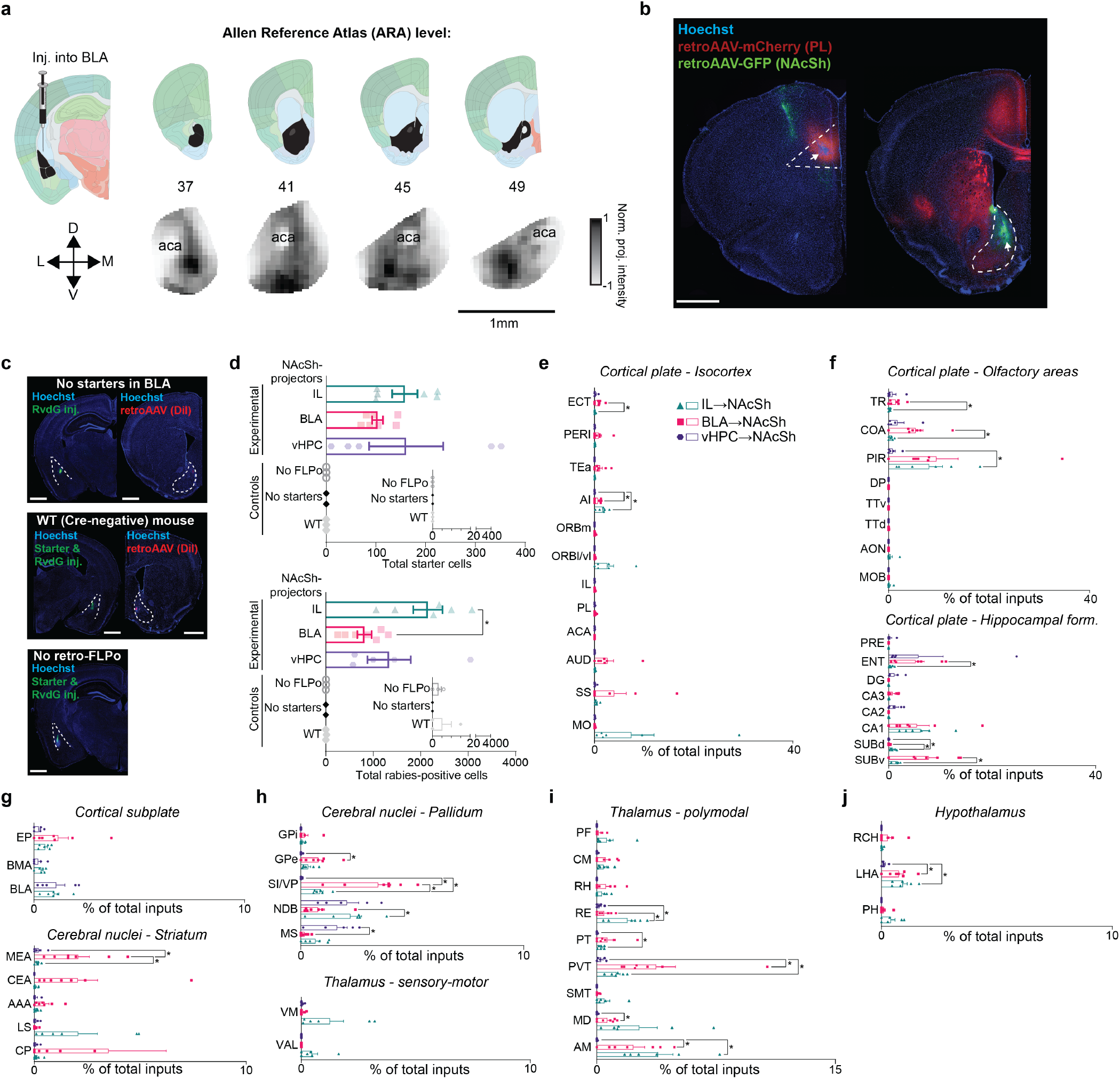
Additional rabies-tracing analyses. (**a**) Projection intensity maps derived from Allen Connectivity Data (Oh et al. 2014) using Brain Street View (https://github.com/Julie-Fabre/brain_street_view) displaying the degree of terminal labeling in NAc following the injection of fluorescent viral tracers into the BLA. Darker colors in heatmap NAc segments (lower row) correspond to more terminal labeling. (**b**) Representative images of injections of retroAAV-mCherry into PL and retroAAV-GFP into NAcSh (for projection overlap quantification in BLA), injection locations indicated by white arrowhead. Scale bar: 1 mm. (**c**) Representative images displaying BLA and NAcSh (if applicable) expression in the three rabies-tracing control groups: no starters (top row), WT mouse injections (middle row), and no retro-FLPo in NAcSh (bottom row). Scale bar: 1 mm. (**d**) Starter cells (top) and the total number of rabies-positive cells (bottom) were distributed across pathways (vmPFC^→^NAcSh pathway slightly enriched in rabies-positive cells vs. BLA^→^NAcSh, p<0.006), with nearly zero labeling observed in either of the 3 control conditions. (**e**) Rabies-positive cells (as a % of total rabies-positive inputs) in regions of the cortical plate (isocortex). More inputs from AI were received by BLA^→^NAcSh neurons (BLA^→^NAcSh vs vHPC^→^NAcSh, p<0.05) and vmPFC^→^NAcSh neurons (vmPFC^→^NAcSh vs vHPC^→^NAcSh, p<0.01). ECT inputs were more numerous to BLA^→^NAcSh neurons (BLA^→^NAcSh vs vmPFC^→^NAcSh, p<0.05). (**f**) Inputs from TR and COA were stronger to BLA^→^NAcSh (BLA^→^NAcSh vs vmPFC^→^NAcSh, both p<0.05). PIR inputs to vmPFC^→^NAcSh neurons were more numerous as compared to vHPC^→^NAcSh neurons (p<0.05). (**g**) Cortical subplate counts did not differ across pathways (top). MEA neurons projecting to BLA^→^NAcSh were more numerous than each other pathway (both p<0.05). (**h**) Inputs to BLA^→^NAcSh from GPe and VP/SI were more numerous, while MS inputs were less numerous, as compared to vHPC^→^NAcSh neurons (all p<0.05). As compared to vmPFC^→^NAcSh neurons, inputs to BLA^→^NAcSh were more numerous from VP/SI, but less numerous from NDB (both p<0.05). vmPFC^→^NAcSh neurons received more input from VP/SI as compared to vHPC^→^NAcSh neurons (p<0.005) (top). Sensory-motor thalamus inputs did not differ across pathways (bottom). (**i**) Inputs to BLA^→^NAcSh neurons from PVT, MD, and AM were more numerous as compared to vHPC^→^NAcSh neurons (all p<0.05), while inputs to vmPFC^→^NAcSh neurons from RE, PVT, PT, and AM were also more numerous relative to vHPC^→^NAcSh neurons (all p<0.05). Inputs from RE to vmPFC^→^NAcSh were also more numerous as compared to BLA^→^NAcSh neurons (p<0.05). (**j**) Within the hypothalamus, inputs to BLA^→^NAcSh and vmPFC^→^NAcSh neurons were more numerous as compared to vHPC^→^NAcSh neurons (both p<0.05). Data mean ± SEM. All analyses were unpaired t-tests with Welch correction. For full reporting of statistics, see **Supplementary Table 1**. Abbreviations (based on Allen Brain Atlas definitions) by panel: (**e**) MO: somatomotor areas, SS: somatosensory areas, AUD: auditory areas, ACA: anterior cingulate area, PL: prelimbic area, IL: infralimbic area, ORBl/vl: orbital area lateral part and ventrolateral part, ORBm: orbital area medial part, AI: agranular insular area, TEa: temporal association areas, PERI: perirhinal area, ECT: ectorhinal area. (**f**) MOB: main olfactory bulb, AON: accessory olfactory nucleus, TTd: taenia tecta dorsal part, TTa: taenia tecta ventral part, DP: dorsal peduncular area, PIR: piriform area, COA: cortical amygdalar area, TR: postpiriform transition area (top). SUBv: subiculum ventral part, SUBd: subiculum dorsal part, CA1: field CA1, CA2: field CA2, CA3: field CA3, DG: dentate gyrus, ENT: entorhinal area, PRE: presubiculum (bottom). (**g**) BLA: basolateral amygdalar nucleus, BMA: basomedial amygdalar nucleus, EP: endopiriform nucleus (top). CP: caudoputamen, LS: lateral septal nucleus, AAA: anterior amygdalar area, CEA: central amygdalar nucleus, MEA: medial amygdalar nucleus (bottom). (**h**) MS: medial septal nucleus, NDB: diagonal band nucleus, SI/VP: substantia innominata/ventral pallidum, GPe: globus pallidus external segment, GPi: globus pallidus internal segment (top). VAL: ventral anterior-lateral complex of the thalamus, VM: ventral medial nucleus of the thalamus (bottom). (**i**) AM: anteromedial nucleus, MD: mediodorsal nucleus of thalamus, SMT: submedial nucleus of the thalamus, PVT: paraventricular nucleus of the thalamus, PT: parataenial nucleus, RE: nucleus of reuniens, RH: rhomboid nucleus, CM: central medial nucleus of the thalamus, PF: parafascicular nucleus. (**j**) PH: posterior hypothalamic nucleus, LHA: lateral hypothalamic area, RCH: retrochiasmatic area. Syringe (**a**, 10.5281/zenodo.4152947) graphic adapted from scidraw.io, coronal mouse brain sections from Allen Mouse Brain Atlas (https://mouse.brain-map.org/static/atlas).

**Extended Data Figure 7:**
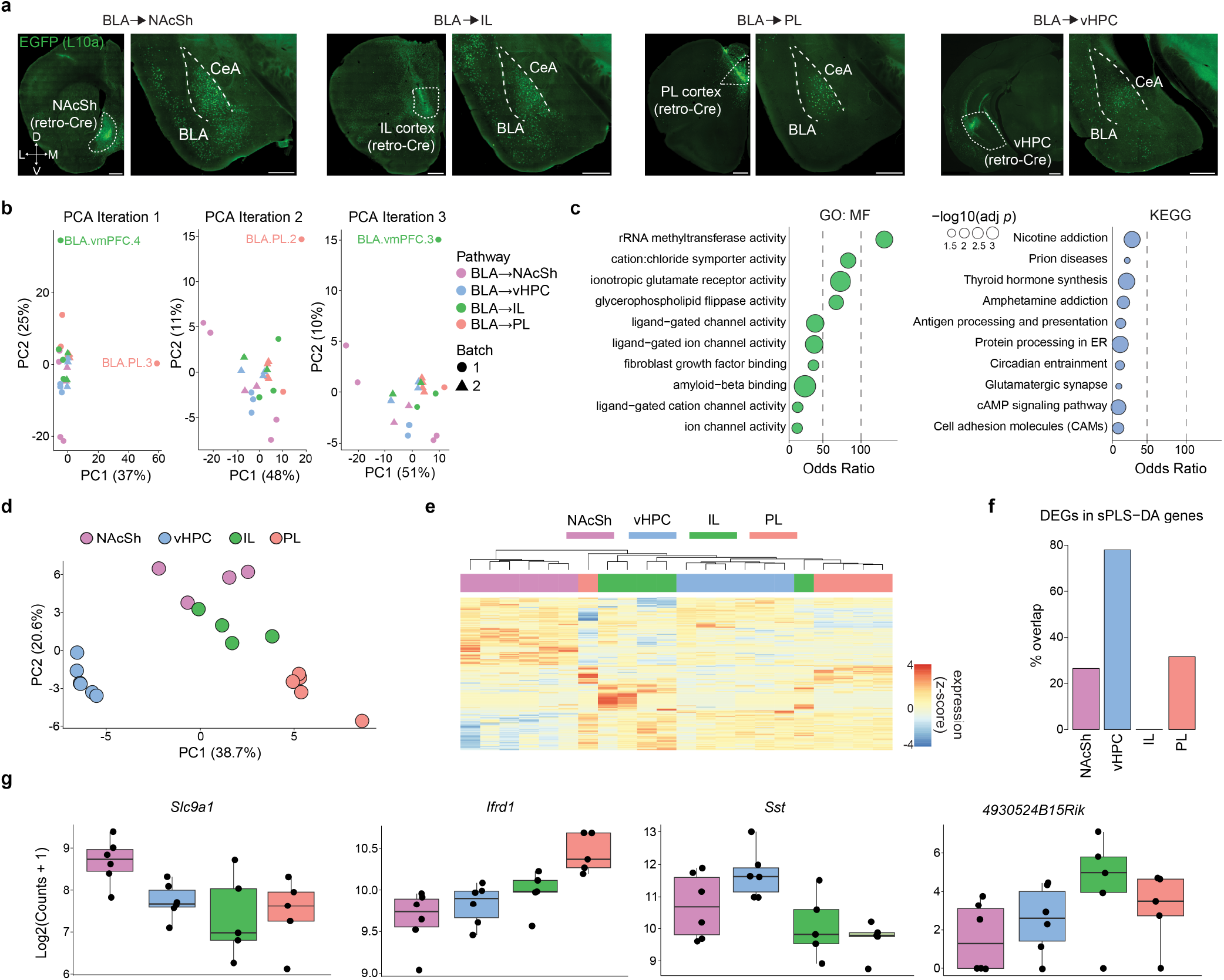
Additional TRAPseq analyses. (**a**) Images depicting retro-Cre injection sites in NAcSh, IL, PL, and vHPC, and EGFP expression in BLA (region punched for sequencing; Scale bars: 500 µm). (**b**) Outliers were identified through three iterations of dimensionality reduction and Mahalanobis distance analysis, resulting in the exclusion of four samples. (**c**) Gene Ontology and KEGG pathway enrichment analysis of genes upregulated in NAcSh highlights enriched molecular functions. (**d**) PCA based on one-vs-all sparse Partial Least Squares Discriminant Analysis (sPLS-DA) reveals region-specific clustering across all BLA projection groups. (**e**) Heatmap displaying the union of top marker genes identified by one-vs-all sPLS-DA for each projection group. (**f**) Proportion of sPLS-DA marker genes that overlap with differentially expressed genes (DEGs) identified by one-vs-all DESeq2 analysis, shown per projection group. (**g**) Expression of representative projection-specific marker genes identified by sPLS-DA, shown as log2(normalized counts + 1).

**Extended Data Figure 8:**
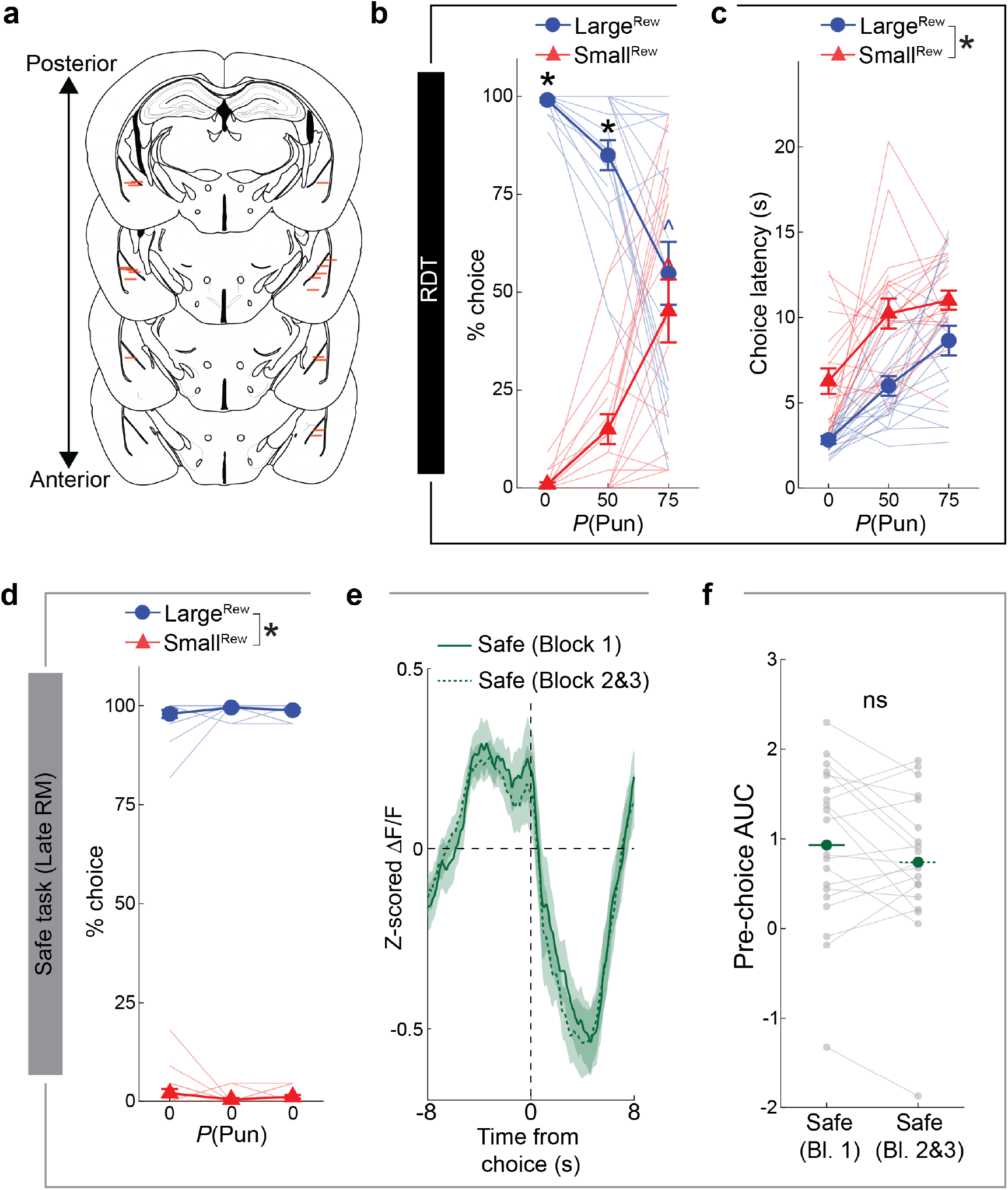
Behavior and photometry measures during RDT and late RM training. (**a**) Ventral extent of photometry optic fiber placement (red lines) in BLA. (**b**) During the fiber photometry RDT recording session, mice (n=20) made fewer Large^Rew^ and more Small^Rew^ choices as punishment risk increased across blocks (within Trial Block effect for Large^Rew^ and Small^Rew^: 75% block versus safe, p<0.001; Large^Rew^ vs Small^Rew^: p<0.001 for 0% and 50% blocks; ANOVA interaction, F(2,36)=25.01, p<0.001). (**c**) Longer choice latencies for Large^Rew^ choices as compared to the Small^Rew^ (F(1,18)=37.48, p<0.0001, repeated measures ANOVA, main effect of Choice Type). (**d**) Large^Rew^ was selected at the near total exclusion of the Small^Rew^ on final RM session before RDT (Late RM) (F(1,19)=15634.89, p<0.0001, repeated measures ANOVA, main effect of Choice Type). (**e**) No difference in choice-aligned photometry signal during the Late RM task across blocks. (**f**) No change in pre-choice photometry signal (area under the curve, AUC) across safe blocks (split into block (Bl.) 1 versus 2/3 for comparison with RDT data shown in **Fig. 6d**). For full reporting of statistics, see **Supplementary Table 1**. Data mean ± SEM. All multiple comparisons (Tukey’s post-hoc tests) were conducted following significant Trial Block (0, 50, 75%) x Choice Type (Large^Rew &^ Small^Rew^) ANOVA interaction. *p<0.05 versus other choice type (**b-d**). ^p<0.05 versus safe block, color corresponding to choice type (**b**).

**Extended Data Figure 9:**
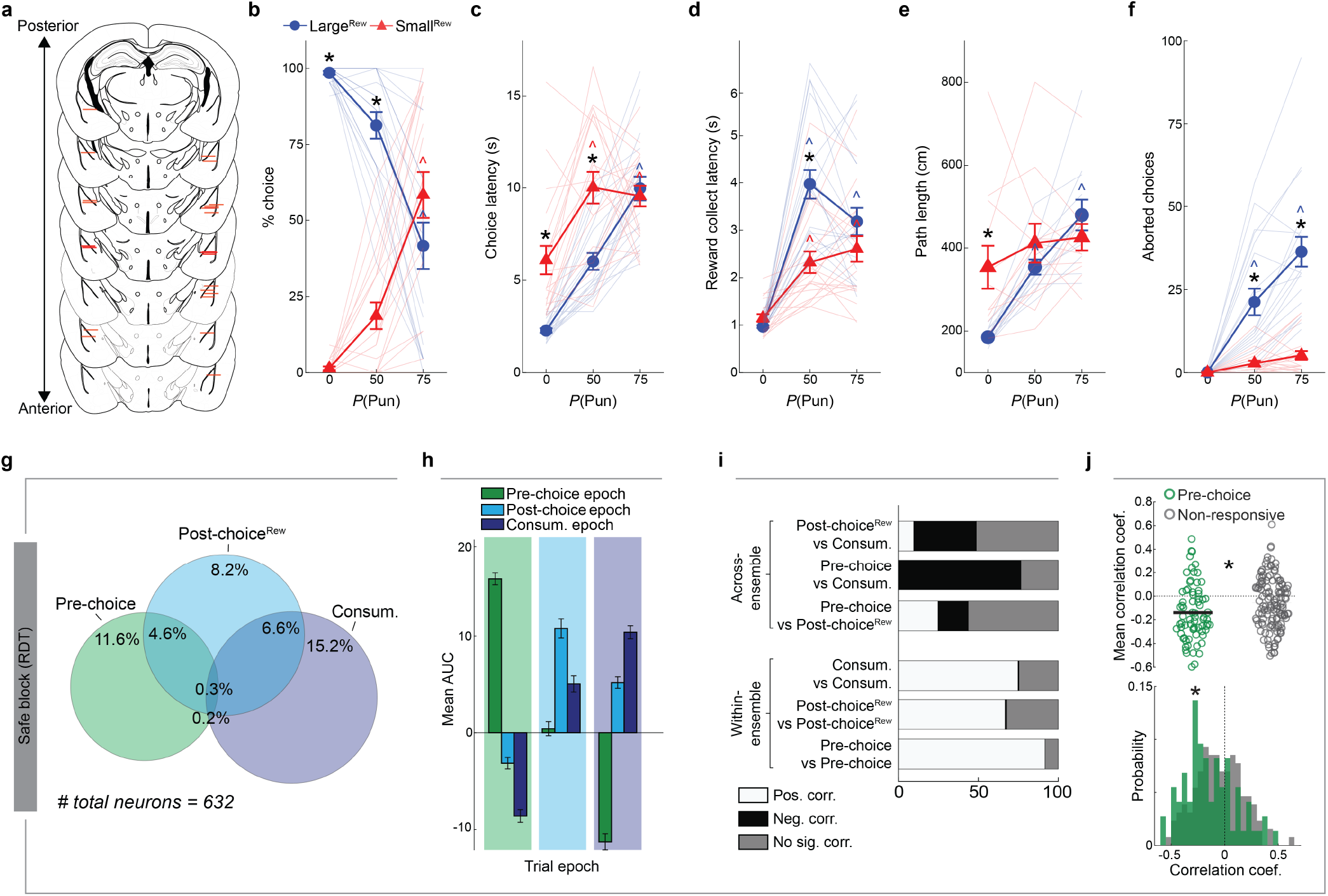
Behavioral and safe ensemble analyses for BLA^→^NAcSh neurons. (**a**) Ventral extent of GRIN lens placement (red lines) in BLA. (**b**) During imaging on RDT, mice (n=20) made fewer Large^Rew^ and more Small^Rew^ choices as punishment risk increased across blocks (within Trial Block effect for Large^Rew^ and Small^Rew^: 75% block versus safe, p<0.001; Large^Rew^ vs Small^Rew^: p<0.001 for 0% and 50% blocks; ANOVA interaction, F(2,36)=25.01, p<0.001). (**c**) Choice latencies were longer for Large^Rew^ choices as compared to the Small^Rew^ (within Trial Block effect for Large^Rew^: 75% block versus safe, p<0.001; within Trial Block effect for Small^Rew^: 50% and 75% blocks versus safe, p<0.001; Large^Rew^ vs Small^Rew^: p<0.001 for 0% and 50% blocks; ANOVA interaction, F(2,36)=10.27, p<0.001). (**d**) Collection latencies were longer for both choice types across risky blocks (within Trial Block effect for Large^Rew^ and Small^Rew^: 50% and 75% block versus safe, p<0.001; Large^Rew^ vs Small^Rew^: p<0.001 for 50% block; ANOVA interaction, F(2,36)=13.55, p<0.001). (**e**) Longer path lengths for Large^Rew^ than Small^Rew^ trials in the safe block (Large^Rew^ vs Small^Rew^: p<0.05 for 0% block; ANOVA interaction, F(2,36)=16.02, p<0.001). (**f**) More aborted choices directed at the Large^Rew^ response-window during risky trial-blocks (Large^Rew^ vs Small^Rew^: p<0.001 for risky blocks), with more Large^Rew^ aborts occurring in these blocks versus the safe block (within Trial Block effect for Large^Rew^: p<0.05 for the 50% and 75% risky blocks versus safe; ANOVA interaction, F(2,18)=34.17, p<0.0001). (**g**) For BLA^→^NAcSh: degree of overlap between Pre-choice, Post-choice^Rew^ and Consum. ensembles. (**h**) For BLA^→^NAcSh: Ensemble AUCs during task epochs (corresponding to **Fig. 6f**). (**i**) For BLA^→^NAcSh: Summary of across and within-ensemble correlations for pairs of ensembles. (**j**) For BLA^→^NAcSh: Higher overall negative correlation for Pre-choice versus non-responsive neurons (p<0.001, unpaired t-test) (top). Negatively-skewed distribution of binned correlation coefficients for Pre-choice (green), relative to non-responsive (grey), neurons (p<0.0001, Kolmogorov-Smirnov test) (bottom). For full reporting of statistics, see **Supplementary Table 1**. Data mean ± SEM. All multiple comparisons (Tukey’s post-hoc tests) were conducted following significant Trial Block (0, 50, 75%) x Choice Type (Large^Rew &^ Small^Rew^) ANOVA interaction. *p<0.05 versus other choice type (**b-f**) or versus non-responsive neurons (**j**). ^p<0.05 versus safe block, color corresponding to choice type (**b**).

**Extended Data Figure 10:**
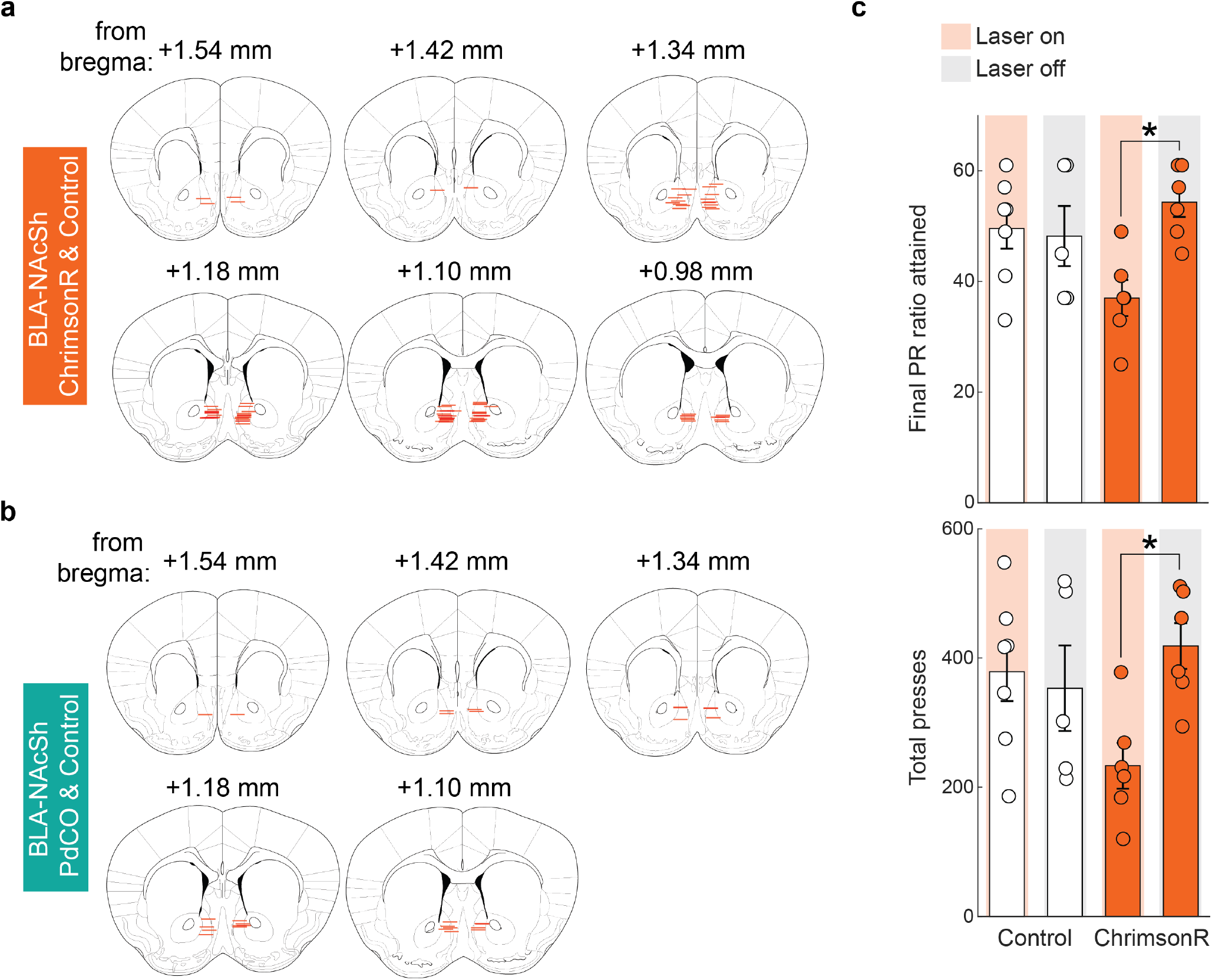
Fiber placement maps and control data for BLA^→^NAcSh optogenetic manipulation. (**a**) Ventral extent of fiber-optic implants for ChrimsonR experiment (red lines) in BLA. (**b**) Ventral extent of fiber-optic implants for PdCO experiment (red lines) in BLA. (**c**) ChrimsonR-mediated BLA^→^NAcSh activation decreased the final PR ratio achieved (top) and the total number of presses made during the PR session (bottom) (ChrimsonR/Laser On vs. ChrimsonR/Laser Off, both p<0.05, ANOVA interaction F-values>5 and p-values<0.05). All multiple comparisons (Tukey’s post-hoc tests) were conducted following significant Treatment (Control & ChrimsonR) x Laser Status (Laser Off & Laser On) ANOVA interaction. For full reporting of statistics, see **Supplementary Table 1**. Data mean ± SEM. *p<0.05

